# Actomyosin-II Proactively Shields Axons of the Central Nervous System from Mild Mechanical Stress

**DOI:** 10.1101/2023.08.09.552549

**Authors:** Xiaorong Pan, Gaowei Lei, Jie Li, Tongshu Luan, Yiqing Hu, Yuanyuan Chu, Yu Feng, Wenrong Zhan, Chunxia Zhao, Frédéric A. Meunier, Yifan Liu, Yi Li, Tong Wang

## Abstract

**Summary:** Pan *et al* found that actomyosin-II-driven radial contractility underpins the resilience of central axons to mild mechanical stress by suppressing the propagation and firing of injurious Ca^2+^ waves. Boosting actomyosin-II activity alleviates axon degeneration in mice with traumatic brain injury.

Traumatic brain injury (TBI) remains a significant and unmet health challenge. However, our understanding of how neurons, particularly their fragile axons, withstand the abrupt mechanical impacts within the central nervous system remains largely unknown. Using a microfluidic device applying discrete levels of transverse forces to axons, we identified the stress levels that most axons could resist and explored their instant responses at nanoscale resolution. Mild stress induces rapid and reversible axon beading, driven by actomyosin-II-dependent radial contraction, which restricts the spreading and bursting of stress-induced Ca^2+^ waves. More severe stress causes irreversible focal swelling and Ca^2+^ overload, ultimately leading to focal axonal swelling and degeneration. Up-regulating actomyosin-II activity prevented the progression of initial injury *in vivo*, protecting commissural axons from degeneration in a mice TBI model. Our study established a scalable axon injury model and uncovered the critical roles of actomyosin-II in shielding neurons against detrimental mechanical stress.

## Introduction

Axons from the white matter of the central nervous system (CNS) are long and thin fibres with parallel orientations, making them susceptible to mechanical insults (Braun et al., 2020; Cavanagh, 1984; Rishal and Fainzilber, 2014; Tang-Schomer et al., 2010). Indeed, sudden mechanical impacts to the head can result in traumatic brain injury (TBI) (Povlishock and Katz, 2005), with diffuse axonal injury (DAI) being the most common pathology (Adams et al., 1989; Johnson et al., 2013), imposing a significant societal burden and impacting individuals, families, and healthcare systems worldwide. The lack of a cure underscores the urgent need for further research and therapeutic advancements. However, it remains unclear how CNS axons manage to withstand mild mechanical stress, which causes 3-5% deformation of the brain tissue during daily activities and contact sports, without incurring injury (Funk et al., 2011; Knutsen et al., 2020). Recent experimental data demonstrate that CNS axons exhibit a remarkable ability to resist both axial stretches (Li et al., 2019; Tang-Schomer et al., 2012; Tang-Schomer et al., 2010) and transverse stress (Gu et al., 2017; Rishal and Fainzilber, 2014), suggesting the existence of intrinsic mechanoprotective mechanisms, acting instantly to defend against the acute mechanical stress. This hypothesized mechanism is likely to stem from the mechanical properties of the axonal cytoskeletal architecture, which consists of a central microtubule (MT) backbone and a peripheral axon cortex organized in periodic rings (Leterrier et al., 2015; Prokop, 2020). Furthermore, recent studies suggest that the mechanical properties of the cortical actin network of the axon may also protect its deeper structures by buffering the mechanical impacts loaded to the axon surface during mild TBI or concussion (Dubey et al., 2020; Kant et al., 2021). The axon cortex is characterized by a membrane-associated periodic cytoskeleton (MPS) that comprises periodically distributed F-actin rings, interlinking spectrin tetramers, non-muscle myosin II (NM-II) motors and other actin-associated proteins (Berger et al., 2018; Costa et al., 2020; D’Este et al., 2015; Leterrier et al., 2015; Vassilopoulos et al., 2019; Wang et al., 2020; Xu et al., 2013; Zhou et al., 2022). While the spectrin-actin lattices provide longitudinal elasticity (Dubey et al., 2020; Krieg et al., 2017), the actomyosin-II, composed of NM-II motor and F-actin, drives radial contraction of the axon cortex (Costa et al., 2020; Wang et al., 2020), maintaining the axon’s surface rigidity (Costa et al., 2020; Smith et al., 2018; Wang et al., 2020). Simulation studies predict that the rigidity of the axon cortex protects against sudden force loading (Dubey et al., 2020; Kant et al., 2021), but whether actomyosin-II is involved in the mechanoprotective response remains unknown.

The influx of Ca^2+^ induced by injury has been proposed to underly the instant axonal response to injury (Wolf et al., 2001), by activating downstream calpains and caspases, ultimately leading to proteolysis of the axon cytoskeleton and initiating the secondary injury (Ma, 2013). Long-range propagation of these destructive Ca^2+^ waves exacerbates the axonal injury by transmitting damage signals to other nearby intact neuronal regions (Mu et al., 2015; Vargas et al., 2015). Whether the transmission of stress-induced Ca^2+^ waves in response to destructive forces can be modulated remains unknown. To answer this critical question, we created an Axon-on-a-Chip (AoC) device (Pan et al., 2022) to investigate how CNS axons can resist scalable mild mechanical stresses by injected medium flow. We found that mild stress triggers a plastic response characterized by rapid and reversible axon beading, accompanied by transient and local Ca^2+^ elevations. The reversible axon beading is mediated by actomyosin-II-driven radial contraction of the transversally stressed axon, which also compresses and stalls the axon organelle trafficking. Importantly, the beading of the stressed axon also inhibited the long-range propagation of the stress-induced Ca^2+^ waves. Down-regulating actomyosin-II activity abolishes the axon beading, worsening the flux-induced axonal injury and ensuing degeneration. Up-regulating actomyosin-II activity leads to a protective effect *in vitro* and a mouse model of mild TBI (mTBI). This work reveals the novel mechanoprotective function of periodic actomyosin-II in enabling the thin axon fibre to withstand mild mechanical challenges.

## Results

### The Axon-on-a-Chip (AoC) captures the instant resilient responses of axons to mild mechanical stress

To simulate the effects of tolerable mild mechanical stress on white matter axons, as seen in mild traumatic brain injury (TBI) or concussion (Fig. 1A, left), we used an Axon-on-a-Chip (AoC) model to determine the range of transverse mechanical stress that cultured hippocampal neuron axons can withstand (Fig. 1A, right). Rat hippocampal neurons were seeded into the two opposing soma chambers (Fig. 1B), the axons of which had extended into the central injury channel after being cultured for 5-6 days *in vitro* (DIV5-6), crossing axons were labelled with β-Tub III (Fig. 1C). To marker the axonal actin cortical meshes, the neurons were transfected with the F-actin marker Lifeact-GFP. On DIV7-8, varying levels of transverse stress were administered to the neurons by injecting culturing medium at different flow rates into the central injury chamber via a syringe pump (Fig. 1D). The instant subcellular responses of the stressed individual axons were live-imaged using spinning disc microscope before, during and after the flow injection (Fig. 1E and Video 1). Cultured medium injected at higher flow rates (100 and 200 μL/min) caused axotomy or diffuse axonal injury (DAI), whereas low flow rates (20 and 50 μL/min) induced reversible axonal beading the axon (Fig. 1F, see also (Pan et al., 2022)). Using an automatic beads detection method (detailed in ref (Pan et al., 2022)), we quantified the axon beading process elicited by four flow rates (20, 50, 100, 200 μL/min for 180 s) (Fig. 1G). The peaks number of beads for all four flow rates were reached within 60 s of the flux, with increasing flow speed resulting in a significant increase in the number of beads (Fig. 1, G and H). Lower flow rates (20 and 50 µL/min) elicited an axonal beading response with more significant reversibility, suggesting that axons can withstand stress below 0.467 Pa, elicited by 50 μL/min flow. In addition, we also noticed that the formation and recovery rates of axon beading are not affected by different flow rates (Fig. S1, A and B).

**Figure 1.**
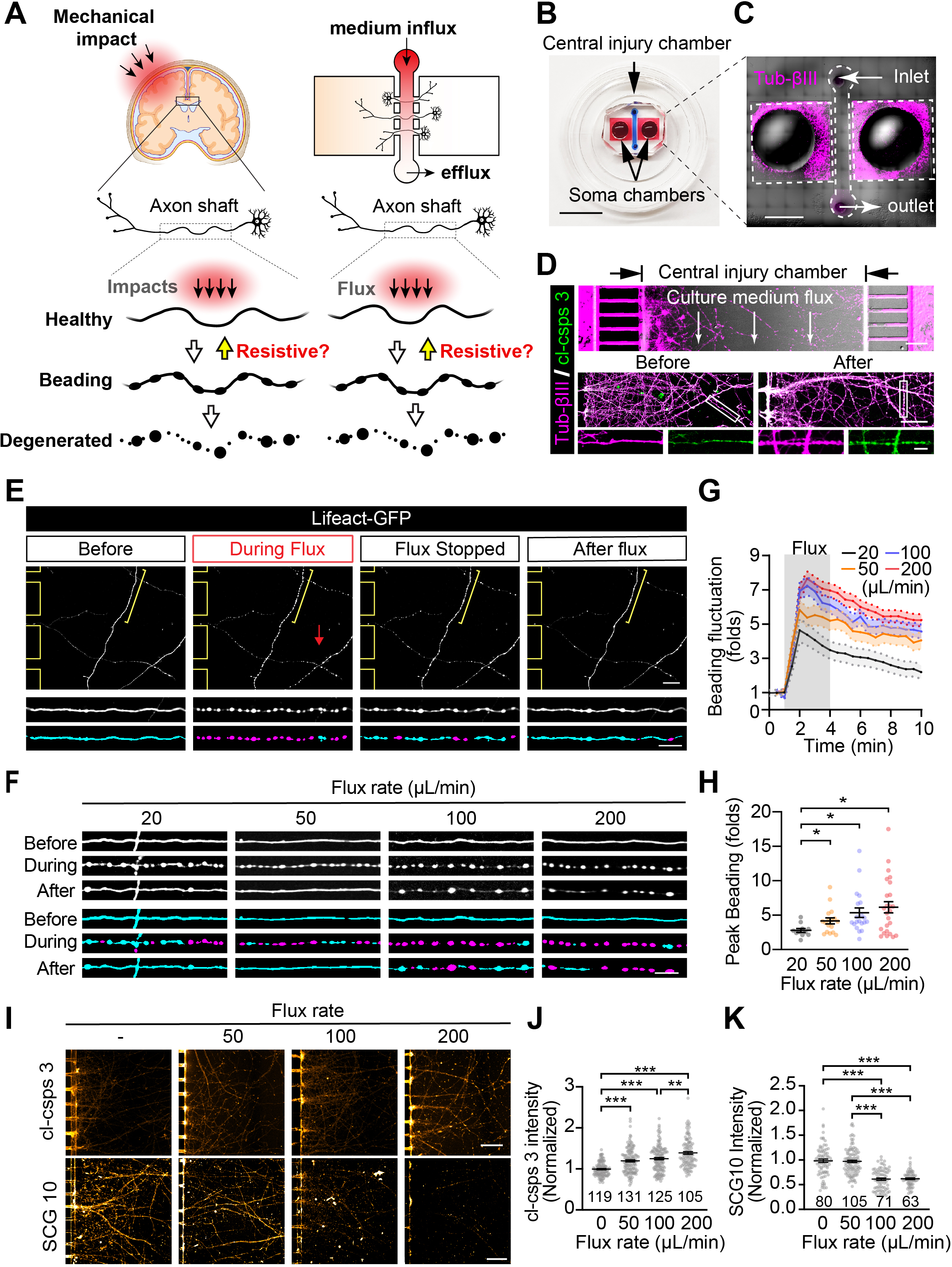
Identify the resistive response of the CNS axons to mild mechanical stress in live neurons cultured in the AoC device. **(a)** Schematic illustration of the axonal resistance hypothesis on the left and the Axon-on-a-Chip (AoC) device on the right. Axonal resistance hypothesis: Mild mechanical impacts to the head can stress the axons of previously healthy neurons, triggering the intrinsic resistance machinery and promoting recovery. However, if the stress exceeds the axons’ resistance capacity, it can cause irreversible axonal injury and eventually lead to degeneration. The AoC device administers different levels of transverse stress to axons by injecting culture medium at varying flow rates, mimicking the stress caused by different levels of impact to the head. This setup allows us to study the instant axonal responses to stress and characterize their biological properties. **(b)** The 3-chambered AoC device is attached to a glass-bottom dish with two opposing soma reservoirs (red) and a single central injury chamber (blue). Bar = 1 cm. **(c)** DIV7-8 mature axons labelled with Tub-βIII (magenta) are mechanically stressed by injecting culture medium into the central injury chamber at different flow rates. Bar = 0.2 cm. **(d)** Axons dual-labelled with Tub-βIII (magenta) and cleaved Caspase 3 (cl-csps 3; green) before (left) and after (right) culture medium flux. Axons highlighted by a white boxed outline are shown magnified with the two fluorescent channels separated in the panels below. Bar = 100 µm (top), 50 µm (middle), and 10 µm (bottom). **(e)** Representative fields showing the axons of the central injury chamber before, during, and after flux with automatically detected “beading” regions (magenta) and regions situated “between” these beads (cyan). The red arrow indicates the flux direction. Bar = 20 µm (top) and 10 µm (bottom). **(f)** Representative time-lapse images show axonal deformation before, during and after different flux rates (mild flux at 20 and 50 µL/min and high flux at 100 and 200 µL/min). “Beading” regions (magenta) and “between” regions (cyan) are shown. Bar = 10 µm. Quantification of the fold changes in **(g)** axon beading and **(h)** peak number of beads in response to different flow-rated flux. N = 25, 20, 17, 11, respectively. **(i)** Representative images of axonal degeneration markers cleaved Caspase 3 and SCG10 following injected flux at the indicated flow rates (non-fluxed control condition indicated by ‘–’). Bar =100 µm. Quantification of **(j)** cleaved Caspase 3 (cl-csps-3) (N = 119, 131, 125, 105, respectively) and **(k)** SCG10 (N=80, 105, 71, 63, respectively) intensity following the injection of flux at the indicated flow rates, with N values shown on the graph. Data represent mean ± s.e.m.; unpaired two-tailed student’s *t-*test; \**p*<0.05, \*\**p*<0.01, \*\*\**p*<0.001.

We next examined the effects of mechanical stress on axonal degeneration and found that higher flow rates (100 and 200 µL/min) caused axonal degeneration of greater severity, which was evidenced by altered immunofluorescent (IF) staining of two axon degeneration markers, cleaved Caspase 3 and SCG10 (Shin et al., 2014), within stressed axons (Fig. 1I). Specifically, as the flow rate increased, the IF staining signals of cleaved Caspase 3 increased (Fig. 1, I, top panels and J), whilst those of SCG10 decreased gradually (Fig. 1, I, bottom panels and K). These findings indicate that axonal beading is a plastic response to mechanical stress and that most axons can withstand transverse stresses below 0.467 Pa (<50 µL/min in our AoC model). However, flux-delivered stresses beyond this range result in irreversible axonal injury and degeneration. The current AoC is, therefore, suitable for investigating mild mechanical stress-induced axonal responses in cultured CNS neurons.

### Radial contraction drives the rapid and reversible beading of stressed axons

Next, to investigate the mechanisms underlying the rapid axonal beading response elicited by the tolerable mechanical stress, we analyzed their dynamic morphological changes induced by a low-speed flux. To label the morphology of the axon, DIV8 hippocampal neurons expressing Lifeact-GFP were stressed by 20 μL/min flux for 180 s. We observed that within the first 5 s, axon beading was induced in the stressed axonal regions, which was largely restored 5-6 min after the withdrawal of stress (Fig. 2, A and B). Based on the circularity of the axonal segments (detailed in ref (Pan et al., 2022)), we separated time-lapse images of the fluxed axons into “beading” (magenta) and non-beading “between” (cyan) regions, in addition to the “total” area (“beading” and “between” combined) of the axon shafts (Fig. 2C). We noticed that immediately following the onset of the flow, the “beading” area increased in size, while both the “between” and “total” area of the axonal shafts were reduced (Fig. 2C), with the total axon area reaching a minimum value (96.03 ± 1.446% of the original) during the 180 s-flux (Fig. 2D). These results indicate that axon beading is not generated by the delayed focal dilation of beading regions but rather by the dramatic thinning of the regions between beads.

**Figure 2.**
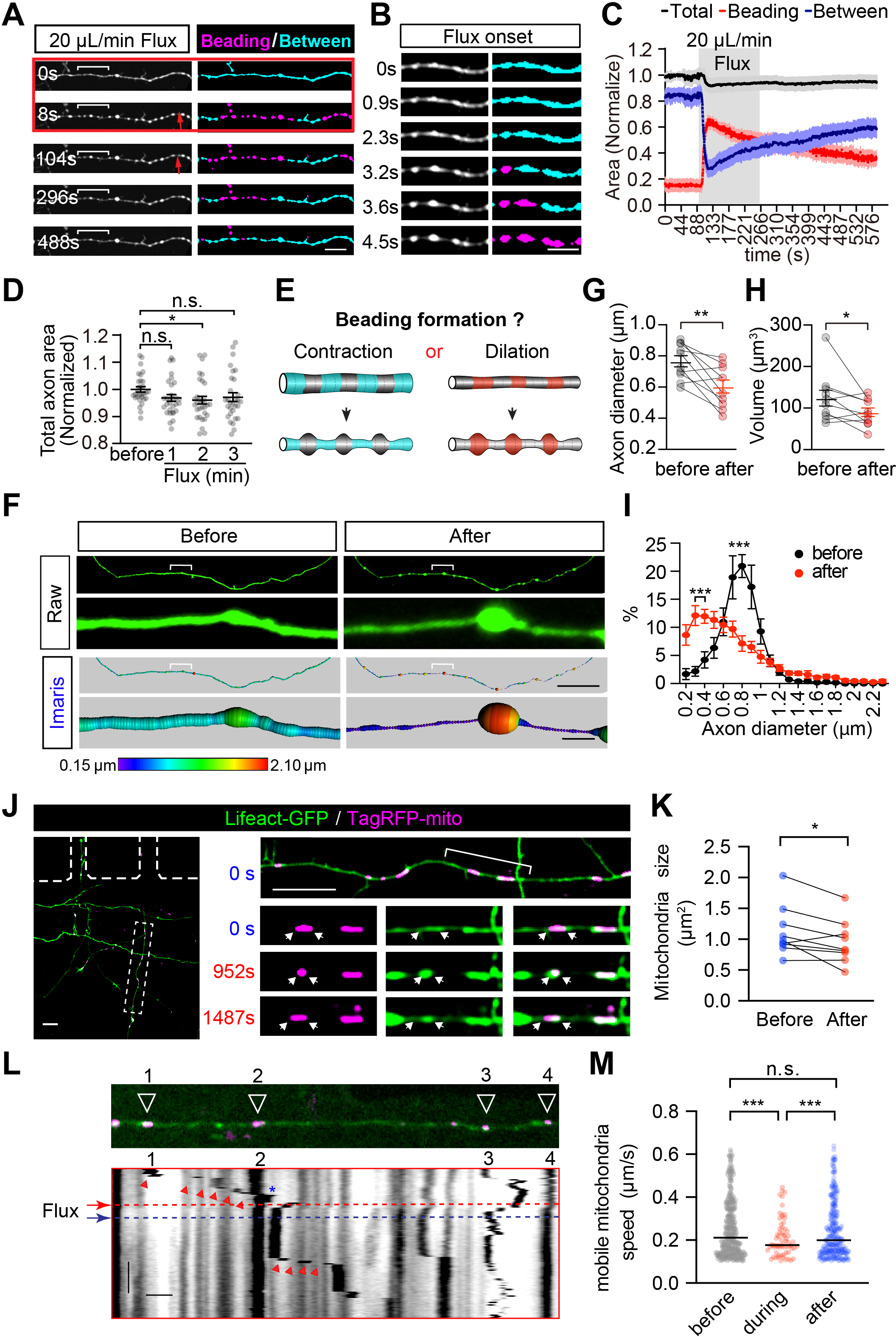
Mechanical stress-induced axon beading results from the radial contraction of stressed axons. **(a)** Rat hippocampal neurons in an AoC device were live-imaged to capture axon beading and morphology changes induced by mild mechanical stress (20 µL/min flux, 180 s). Beading axonal segments are shown in magenta, and non-beading segments between the beads are shown in cyan. The direction of the injected flow is indicated with a red arrow. Bar = 10 μm. **(b)** Magnification of the bracketed region shown in **(a)** depicts the rapid beading process that occurs during the initial 4.54 s following the onset of flux. Bar = 5 μm. **(c)** Curves showing the instantaneous changes in beading (red), between (blue) and total (black) axonal areas during the 20 µL/min flux. **(d)** Quantification of the total axonal area during the 20 µL/min flux. N = 30. **(e)** Models depicting two possible mechanisms that could drive axon beading. (Left) axonal regions between the beads could contract to form the “string” (blue), or (right) the beaded regions themselves could dilate to form the “beads” (red). **(f)** The z-stack raw images show morphological changes in the same axon before and after a 50 µL/min flux, with 3D Imaris renderings shown at the bottom. Axonal diameter is colour-coded. Bar = 30 μm (top) and 2 μm (bottom). **(g-i)** Paired analysis of the Imaris rendered 3D surfaces shows flux-induced morphological changes within the same axon, with quantification of **(g)** the median axon diameter, **(h)** axonal volume, and **(i)** the distribution of axonal diameter. N = 10. **(j)** Time-lapse images show the instant shape changes of inner mitochondria induced by the low-speed flux (50 μL/min, 180 s). The boxed region is amplified in the right panels. Shape changes of the mitochondria are indicated with arrowheads. Bar=10 μm. **(k)** Paired comparison of mitochondria size fluctuations induced by low-speed flux. N=9. **(l)** Axonal trafficking of the intra-axonal mitochondria during the 50 μL/min flux. The perpendicular lines in the kymograph indicate the pausing stage of the mitochondria, whereas the tilted slopes indicated by arrowheads are the mobile stage. x-axis Bar = 10 μm; y-axis Bar = 500 s. **(m)** Quantification of the average speed of the mobile mitochondrial trajectories of **(l)**. N = 328, 67, 275, respectively. Data represent mean ± s.e.m.; (d) unpaired two-tailed student’s *t*-test; (g-i, k) paired two-tailed student’s *t*-test; (m) Welch’s *t*-test; \**p*<0.05, \*\**p*<0.01, \*\*\**p*<0.001; n.s. Non-significant.

Within 5 s, the transformation of the axon from a cylinder shape to a “string of beads” shape had occurred – likely generated by the contraction of the area between the beads, forming the thin “strings” that are observed between “beads” (Fig. 2E). To explore this transformation further, 3D z-stack images of live axons immediately before and after the administration of mild stress (50 μL/min flux for 180 s) were analyzed using Imaris software to extract volume parameters for use in paired comparison (Fig. 2F and Video 2). The axonal diameter in 8 out of 10 stressed axons was significantly reduced, with the median diameter decreasing from 0.765 ± 0.036 μm to 0.604 ± 0.042 μm (Fig. 2G). There was an overall reduction in axonal volume (decreased to 79.39 ± 9.432 %) (Fig. 2H), immediately after the flux. The main peak of the diameter profile of these stressed axons demonstrated a significant left shift (Fig. 2I), suggesting the diameter reduction. Next, we used time-lapse Structured Illumination Microscopy (SIM) to capture these morphological changes with sub-diffraction resolution. By doing so, we identified the continuous reduction in the mean diameter of most axons during imaging and noted the formation of beads during stress, with partial recovery after cessation of stress (Fig. S2A and Video 3). Diameter distribution analysis revealed that axons could withstand mechanical stress and partially recover over time (Fig. S2, B-E). Together, the 2D and 3D time-lapse data demonstrate that the acute radial contraction of the stressed axonal cortex drives the observed axon beading.

Following this, we investigated whether the distribution of stress-induced axonal beads is related to the presence of large organelles, which can significantly expand the diameter of the corresponding axonal region (Wang et al., 2020). Axons of hippocampal neurons expressing TagRFP-mito were subjected to low-speed flux, and we found that the distribution of mitochondria was highly correlated with that of stress-induced axonal beads (Fig. S2F). The ratio of mitochondria existing in the beading area was significantly increased after the flux (Fig. S2, G and H). Conversely, the axonal regions lacking mitochondria underwent a more dramatic radial contraction, forming a “string” in the stressed axon between the axonal beads (Fig. S2G, asterisks). Of note, the distribution of these axonal beads is not periodic, with intervals showing no obvious auto-correlation (Fig. S2, I and J). These data, therefore, revealed that some axonal beads may be caused by the presence of inner organelles such as mitochondria.

Since radial contractility of the axon cortex can significantly affect the mobility of large axonal organelles (Wang et al., 2020), we investigated whether the shape and dynamics of axonal mitochondria were altered during the flux and found that axonal mitochondria were indeed compressed and shortened during the flux (Fig. 2, J and K; and Video 4), with kinetics highly consistent with the shape change of the contracted axonal membrane outside the indicated organelle (Fig. 2J and Video 4). Additionally, the trafficking of previously mobile mitochondria was paused during and immediately after the flux (Fig. 2L, bottom; and Video 5), and some of the paused mitochondria restored their mobility several minutes later (Fig. 2L and Video 5), suggesting the reversibility of the axonal radial contraction. Further analysis using automated tracking showed a significant reduction in the speed of mitochondrial trafficking during flux (Fig. 2M and Video 5), indicating that mild stress triggers the reversible radial contraction of the axonal cortex, which temporarily compresses and halts previously mobile mitochondria. These results indicate that the axon’s typical “string of beads” shape is generated by the rapid and reversible contraction of the axonal cortex, which significantly impacts the morphology and trafficking of axonal organelles.

### Actomyosin-II mediates the reversible axon beading induced by mild mechanical stress

To determine the impact of mild mechanical stress on the axonal surface, we used Scanning Electron Microscopy (SEM) to examine their surface integrity. We found that although the axons thinned significantly (Fig. 3A) and exhibited significantly more fluctuations in diameter (Fig. 3B), the integrity of the axon surface remained intact. Axon radial contractility is driven by actomyosin-II, a complex comprising molecular motor NM-II and F-actin rings (Costa et al., 2020). Accordingly, we examined whether actomyosin-II controls the stress-induced contraction of the axonal cortex. Using time-lapse SIM, we observed that the periodic actomyosin rings labelled by Lifeact-GFP underwent dramatic shortening between the beads (Fig. 3, C, white arrowheads, D and E; and Video 6). During the reversal of the beading, the length of these actomyosin rings partially recovered (Fig. 3, C, white arrowheads, D and E; and Video 6). As previously reported (Berger et al., 2018; Wang et al., 2020), activated NM-II are assembled into unipolar or bipolar filaments. By using antibodies that recognize the C-terminal (αCT) and N-terminal (αNT) regions of NM-IIB (Fig. S3A) and performing 3D-Stimulated Emission Depletion Microscopy (3D-STED), we found that assembled NM-IIB filaments are abundant in the shafts of mature axons, with most being unipolar (Fig. S3, B, arrowheads and C, asterisks), with bipolar NM-II filaments rarely occurred (Fig. S3D). These findings suggest that activated actomyosin-II filaments are abundantly distributed along the shafts to drive the contraction of the axon cortex.

**Figure 3.**
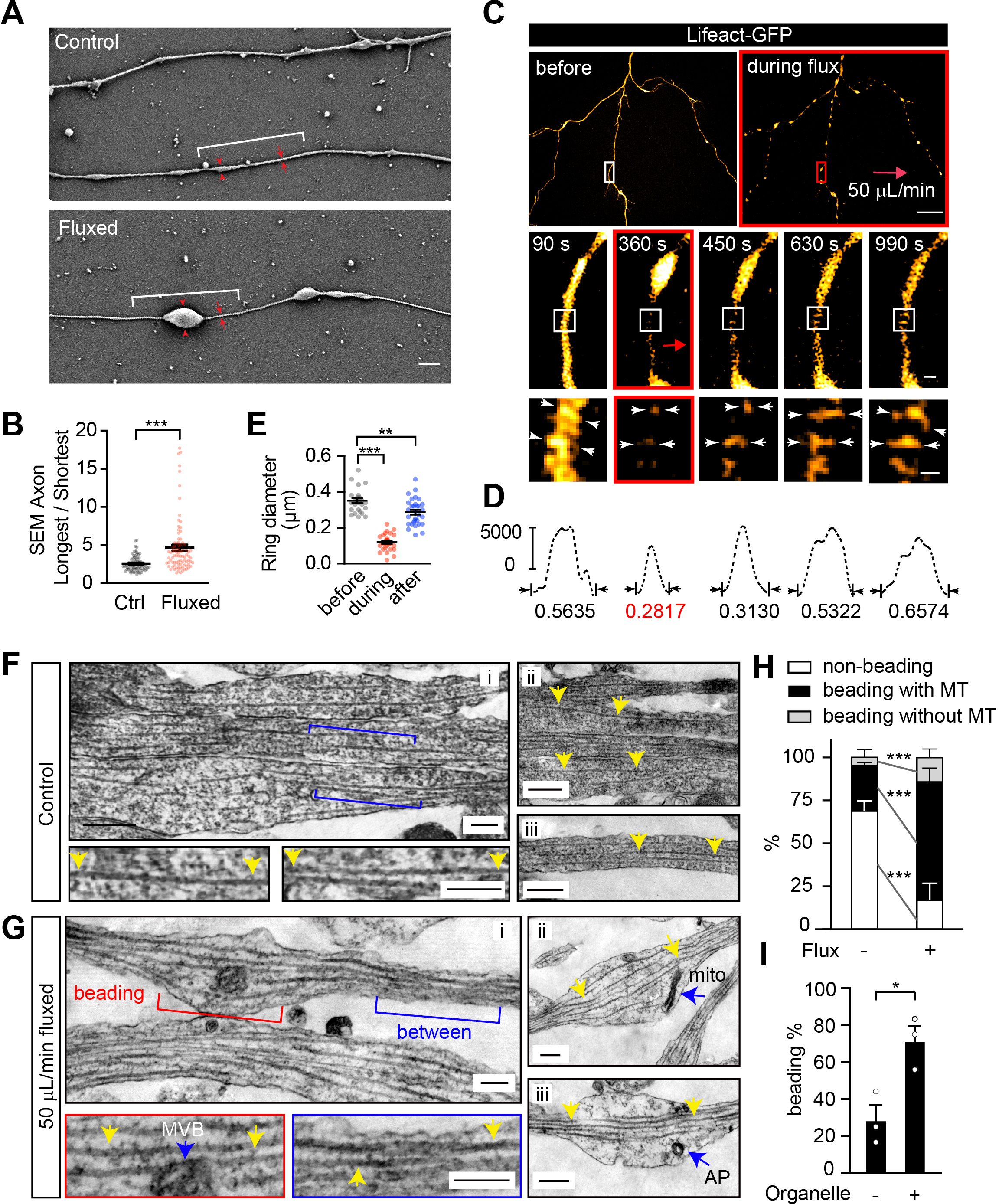
Radial contraction of actomyosin rings underlies flux-induced axon beading. **(a)** Representative SEM images illustrate axon beading induced by low-speed (50 μL/min) flux. Red arrows indicate the fluctuation of axon diameter. Bar = 1 μm. **(b)** Quantification of axon diameter fluctuations, calculated by the longest to shortest diameters per 10 μm length of the axon, in control (Ctrl) and fluxed neurons. N = 80, 86, respectively. **(c)** Representative time-lapse SIM images of axons expressing Lifeact-GFP, with actin rings magnified in the bottom panels showing the dynamic diameter change of actin rings before, during, and after the flux, with white arrowheads pointing to the diameter change and red arrow indicating the flux direction. Bar = 10 μm (top), 0.5 μm (middle) and 0.2 μm (bottom). **(d)** The diameter changes of indicated actin rings in **(c)**. y-axis indicates the fluorescent units. **(e)** Quantification of the actin ring diameter during the 50 μL/min flux. N = 26, 29, 31, respectively. **(f-g)** Representative TEM images show the cytoskeletal change in **(f)** control and **(g)** 50 μL/min fluxed axons. Intact microtubule (MT) tracks are highlighted by yellow arrows, and organelles (multivesicular body, MVB; autophagosome, AP; mitochondria, mito) are marked by blue arrows. Bar = 200 nm. **(h)** Quantification of the percentage of axonal segments with either non-beading, beading with MTs, or beading without MTs, in control and fluxed axons. N = 3 preparations. **(i)** Quantification of the percentage of axon beads containing organelles. N = 3 preparations. Data represent mean ± s.e.m.; unpaired two-tailed unpaired *t-*test; \**p*<0.05, \*\**p*<0.01, \*\*\**p*<0.001.

Since microtubule (MT) tracks are crucial for the formation of focal axonal swellings (FAS) – an axonal beading structure that marks irreversible axonal injury (Datar et al., 2019; Gu et al., 2017; Prokop, 2021; Qu et al., 2017), we explored the integrity of axonal MT bundles in beading axons using Transmission Electron Microscopy (TEM). We found that MTs are oriented in parallel tracks in non-fluxed axons (Fig. 3F, yellow arrowheads). After mild flux, the beading area significantly increased (Fig. 3, G and H). However, in most of these beads, the MT tracks remained intact (Fig. 3, G, yellow arrowheads and H, black bars). Additionally, after the flux, most of the axonal beads contained organelles (Fig. 3G and I), verifying that the presence of organelles in the axon may contribute to the formation of the beads. We further investigated the potential role of MTs in axon beading by treating neurons with MT polymerization inhibitor Nocodazole and MT stabilizer Taxol. However, neither Nocodazole nor Taxol treatment affected the kinetics of axon beading (Fig. S3E), as no changes in the curve of flux-induced axon beading (Fig. S3F), peak beading density (Fig. S3G) or axon volume (Fig. S3H) were observed. These data, therefore, suggest that MT disorganization is unlikely to be the cause of underlying stress-induced reversible axon beading.

Next, we used two pharmacological inhibitors of NM-II activity, Blebbistatin (Kovács et al., 2004) and ML-7 (Kovács et al., 2004), to examine whether flux-induced axon beading is affected by NM-II inactivation. As shown in Fig. 4A-C, and Video 7, pre-incubation of neurons with either Blebbistatin (+BLB; 50 μM) or ML-7 (+ML-7; 10 μM) for 30 minutes significantly abolished (+BLB) or reduced (+ML-7) the axon beading induced by mild stress (50 μL/min flux for 180 s). Additionally, when pre-incubating neurons with NM-II activator Calyculin A (+CA; 50 nM), which enhances the consecutive radial contraction of axon shafts (Costa et al., 2020; Wang et al., 2020), significantly decreased the stress-induced axon beading by 50%, suggesting that axonal segments with over-activated NM-II are not sensitive to mild mechanical stress (Fig. 4, A-C and Video 7). Furthermore, upon disruption of F-actin rings using latrunculin B (+LatB; 5 μM, 30 minutes), we found that the stress-induced axonal beading was abolished (Fig. 4, A-C; and Video 7). These data suggest that coordinated actomyosin-II activity is required for stress-induced axon beading.

**Figure 4.**
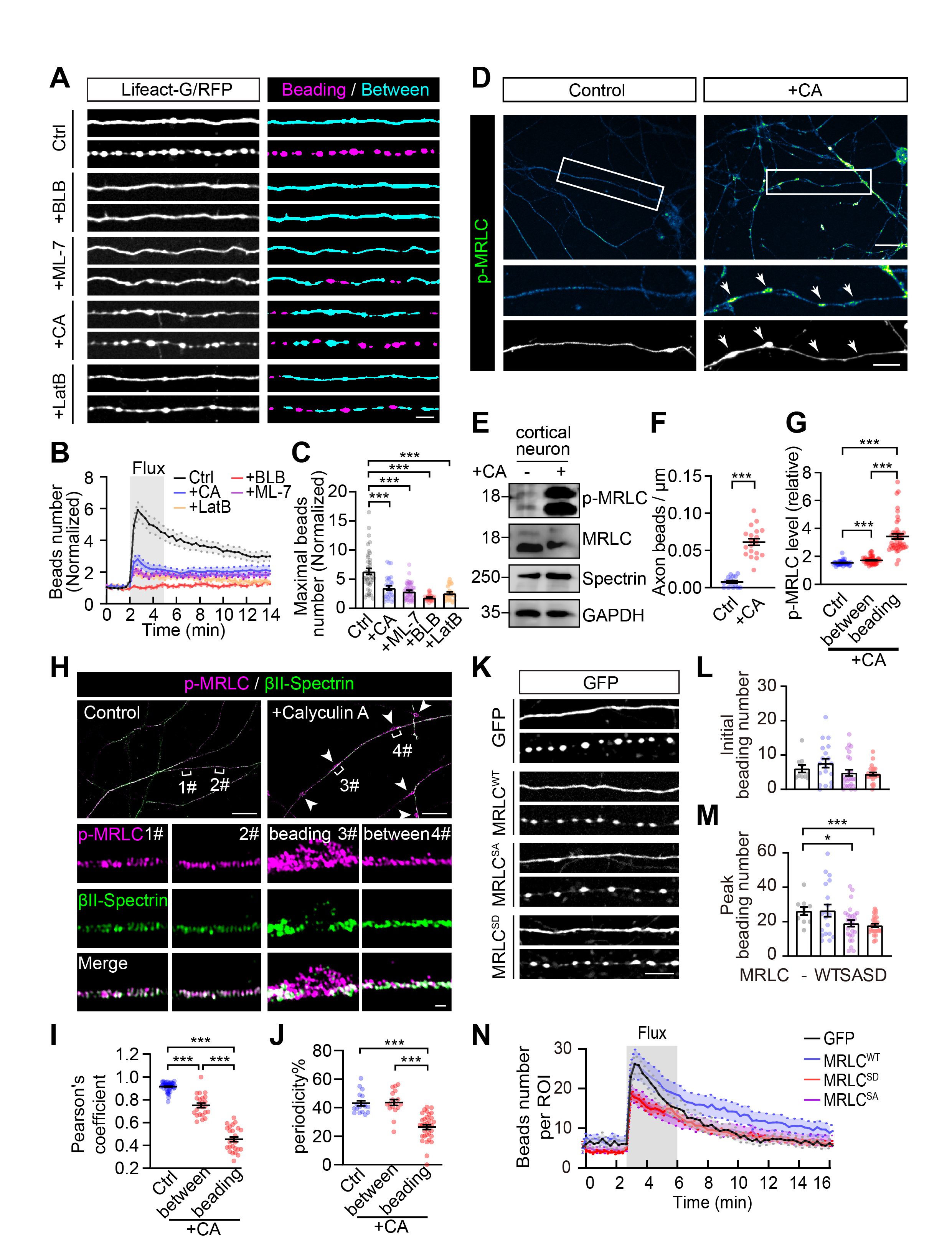
Flux-induced reversible axon beading is driven by actomyosin-II. **(a)** Lifeact-GFP expressing neurons were either untreated (Ctrl) or pre-treated with either Blebbistatin (+BLB; 50 μμ), ML-7 (+ML-7; 10 μμ), Calyculin A (+CA; 50 nM), or Latrunculin B (+LatB; 5 μμ), prior to being stressed with 50 μL/min-flux for 180 s. Representative time-lapse images show the deformation of the same axons before (top) and during (bottom) the flux for each condition, with “beading” regions shown in magenta and “between” regions shown in cyan. Bar = 5 μm. Quantification of **(b)** the number of axon beads formed and **(c)** the peak number of axon beads formed in response to the 50 μL/min flux for 180 s. N = 41, 29, 49, 30, 23, respectively. **(d)** IF staining of phosphorylated MRLC (p-MRLC) after CA treatment. The white boxed outline is shown magnified at the bottom, with the beading annotated by arrows. Bar = 20 μm (top), 10 μm (bottom). **(e)** Western blot showing the level of the p-MRLC, total MRLC, βΙΙ-Spectrin and GAPDH following CA treatment in cultured cortical neurons. **(f)** Quantification of axon beading induced by CA treatment. N = 16, 19, respectively. **(g)** Quantification of the p-MRLC intensity in **(d)**. N = 41, 45, respectively. **(h)** Representative 3D-SIM image showing the localization between periodic actin rings (βII-Spectrin; green) and p-MRLC (magenta). Bracketed regions 1# and 2# in control and beading region #3 and between region #4 in the CA treated axon are shown magnified below. Bar = 10 μm (top) and 0.5 μm (bottom). **(i)** Quantification of the Pearson’s correlation coefficient reflecting the colocalization between actin rings and p-MRLC. N = 55, 25, 23, respectively. **(j)** Quantification of the spacing between periodic actin rings. N = 17, 17, 33, respectively. **(k)** Representative time-lapse images showing axons infected with AAV virus expressing either wild-type MRLC (MRLC^WT^), an inactive MRLC mutant (MRLC^SA^), a constitutively active MRLC mutant (MRLC^SD^) or an empty vector, with the deformation of the same axon induced by 50 μL/min-flux shown below the corresponding condition. Bar = 10 μm. **(l-n)** Quantification of (l) the initial number of beads, **(m)** the peak number of beads, and **(n)** the curves showing the changes in the number of beads per region of interest (ROI) induced by the 50 μL/min-flux. N = 10, 19, 26, 25, respectively. Data represent mean ± s.e.m.; unpaired two-tailed student’s *t*-test; \**p*<0.05, \*\*\**p*<0.001;

Since Calyculin A can increase the phosphorylation of the myosin regulatory light chain (p-MRLC), the regulatory light chain of NM-II, which in turn enhances the consecutive radial contraction of axon shafts (Costa et al., 2020; Wang et al., 2020), we next examined the effect of Calyculin A on axonal beading and found that Calyculin A treatment increased both the p-MRLC level (Fig. 4, D and E) and beading density (Fig. 4F) in axons, with the highest level of p-MRLC accumulating within the beading area (Fig. 4, D, arrowheads and G). We further resolved the periodic distribution of actomyosin-II rings by 3D-SIM of the p-MRLC. By doing so, we found that most of the p-MRLC periodic structure in control axons overlaps with βII-Spectrin (Fig. 4, H and I). In contrast, after Calyculin A treatment, the colocalization between p-MRLC and βII-Spectrin and the periodicity of p-MRLC were significantly reduced, especially in the beading regions (Fig. 4, H, arrowheads, I and J). To further manipulate NM-II activity, we transfected neurons cultured in an AoC device with either the inactive MRLC mutant S19A/T18A (SA), the constitutively active MRLC mutant S19D/T18D (SD) (Beach et al., 2011), or wild-type MRLC (WT). We found that the beading density in non-fluxed axons is unaffected (Fig. 4, K and L). In contrast, under mild stress (50 μL/min, 180 s), the capacity of axon beading in both NM-II inactivated MRLC (SA) and constitutively active MRLC (SD) neurons was significantly reduced (Fig. 4K), reflected by the significantly reduced peak beading densities (Fig. 4M), and altered beading kinetics (Fig. 4N), as compared to MRLC (WT) transfected neurons. These results demonstrate that the coordinated actomyosin-II activity is both necessary and sufficient to drive the radial contraction of the fluxed axon, leading to rapid and reversible axon beading within seconds after the onset of stress.

### Axon beading restricts Ca^2+^ elevation to mechanically stressed regions

Next, we investigated the physiological functions of stress-induced axon beading. In the AoC device, we found that within the same axon, flux-induced beading was restricted to the stressed distal region (Fig. 5, A, denoted by 2# and B; Fig. S4A). In contrast, the morphology of non-fluxed proximal segments in the soma chamber showed no significant change after the flux (50 μL/min for 180s) (Fig. 5, A, denoted by 1# and B; Fig. S4A). In addition, the phenomenon of regionally restricted beading formation was also observed in axons stressed by higher flow speed (200 μL/min for 180s), with the non-fluxed proximal axonal regions remained structurally intact (Fig. S4B). These data suggest that the reversible beading is restricted to the stressed axonal regions.

**Figure 5.**
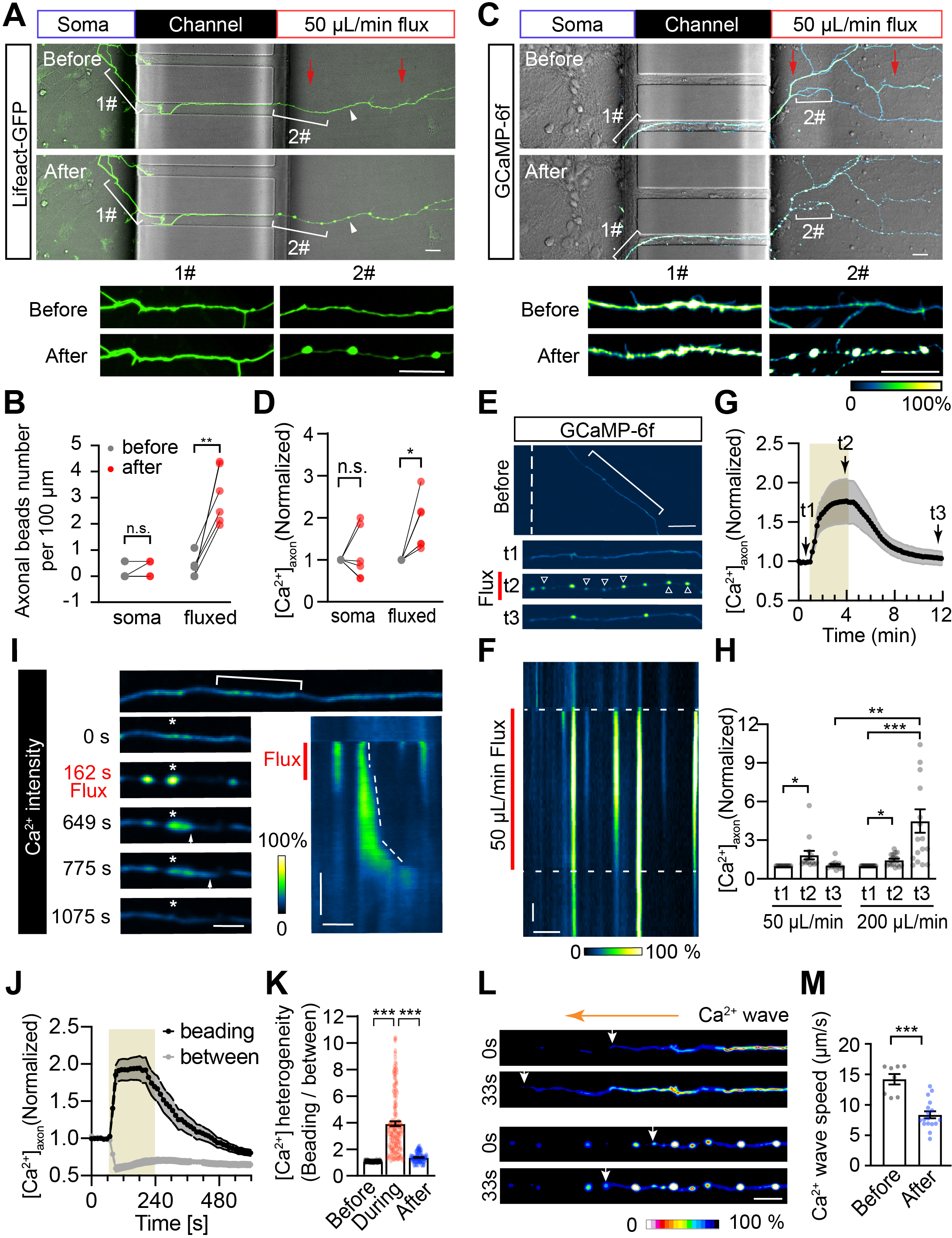
Both stress-induced axonal beading and Ca^2+^ elevation are spatially restricted. **(a)** Representative image of non-stressed axonal segments in the soma chamber (Soma) and stressed segments within the injury chamber (Flux) of the same neuron before and after 50 μL/min-flux for 180 s. Bracketed regions in the soma (1#) and bracketed regions in flux (2#) are shown magnified below. Bar = 20 μm. **(b)** Paired comparison of the number of beads in non-stressed (soma) and stressed segments (fluxed) before and after the flux. N = 6. **(c)** Axons expressing GCaMP-6f were stressed with 50 μL/min-flux for 180 s. [Ca^2+^]_axon_ intensity was colour-coded in the non-stressed (Soma) and the stressed (Flux) axonal segment. Bar = 20 μm. **(d)** Paired comparison of the [Ca^2+^]_axon_ in non-stressed (soma) and stressed segments (fluxed). N = 6. **(e)** Spatial and temporal changes in [Ca^2+^]_axon_ before (t1), during (t2) and after (t3) the 180 s of 50 µL/min flux, marked by the red line. The bracketed region is amplified at the bottom, with arrowheads indicating axon beads. Bar = 10 μm. **(f)** Kymographs of magnified axons in **(e)**, x-axis Bar = 5 μm, y-axis Bar = 20 s. **(g)** Quantification of the total [Ca^2+^]_axon_ fluctuation induced by the 50 μL/min flux. The flux span is indicated with shades, with arrows indicating t1, t2 and t3. **(h)** Quantification of [Ca^2+^]_axon_ at t1, t2 and t3 in 50 or 200 μL/min fluxed groups. N = 14, 17. **(i)** Time-lapse images showing the kinetics of the [Ca^2+^]_axon_ along the beading axons, with the beading region indicated with an asterisk and the [Ca^2+^]_axon_ spreading indicated with an arrowhead. Kymograph is shown on the right, with a dashed line indicating the spreading of the [Ca^2+^]axon. Bar = 20 μm (left), x-axis Bar = 20 μm and y-axis Bar = 200 s (right). **(j)** Quantification of the flux-induced [Ca^2+^]_axon_ fluctuation in “beading” and “between” regions. N = 10. **(k)** Quantification of the heterogeneity of [Ca^2+^]_axon_. N = 115, 150, 110, respectively. **(l)** Comparison of the spreading speeds of the [Ca^2+^]_axon_ waves along the same axon before and after the 50 μL/min-flux, which induced axon beading. Ca^2+^ intensity is colour-coded. Arrows indicate the frontier of the wave. Bar = 5 μm. **(m)** Quantification of **(l)**. N = 8, 17, respectively. Data represent mean ± s.e.m.; In (b, d) paired student’s *t*-test; in (h, k, m) two-tailed student’s *t*-test. \**p*<0.05, \*\**p*<0.01, \*\*\**p*<0.001; n.s. non-significant.

We then explored the mechanisms underlying the restriction of reversible axonal beading to stressed regions. The communication between an injured axon and its soma occurs through the long-range spreading of axonal Ca^2+^ waves, defined as the [Ca^2+^]_axon_, which originates from the injury site and spreads bidirectionally, triggering secondary degenerative responses en route (Rishal and Fainzilber, 2014; Vargas et al., 2015; Witte et al., 2019). In neurons expressing the Ca^2+^ sensor GCaMP-6f, we observed that mild mechanical stress (50 μL/min flux) induced significant elevation of [Ca^2+^]_axon_, which was restricted to the distal region of the axon which was directly exposed to the mechanical stress (Fig. 5C, distal region (2#) and D; Fig. S4C). However, such elevation was absent in the proximal region inside the soma reservoirs (Fig. 5C, proximal region (1#) and D; Fig. S4C). The elevation of the [Ca^2+^]_axon_ coincided with the axon beading process, both of which are regionally restricted to the mechanically stressed axon parts (Fig. S4, D, proximal region (1#) and E), suggesting that stress-induced radial contraction may correlate with the restriction of elevated [Ca^2+^]_axon_.

To explore the relationship between stress-induced axon beading and the resulting [Ca^2+^]_axon_ wave, we examined their kinetics whilst applying mild mechanical stress. We found that compared to the Ca^2+^ level prior to flux onset (t1), the [Ca^2+^]_axon_ was significantly increased by 1.310 ± 0.127 fold in the entire stressed axon, with the peaks achieved at approximately 160 s (t2) and restored around 480 s following flux onset (t3) (Fig. 5, E and G). Interestingly, the Ca^2+^ elevation triggered by mild stress was only prominent in the beading regions, which were restricted and soon reduced to the baseline. Such reversibility in the elevation of [Ca^2+^]_axon_ sharply contrasts with that of the axons subjected to higher flow rates, which instead caused the progressive increase in Ca^2+^ surges that ultimately led to irreversible axonal injury (Fig. 5H). Blocking flux-induced [Ca^2+^]_axon_ elevation using Ca^2+^ chelators EDTA or BAPTA-AM significantly reduced the level of axonal beading (Fig. S5, A-C), suggesting that [Ca^2+^]_axon_ is required to initiate axonal beading. These data indicate that the reversibility of [Ca^2+^]_axon_elevation is highly related to the reversibility of beading.

Because the [Ca^2+^]_axon_ in mildly stressed axons is highly restricted (Fig. 5, E-G), we next examined whether axon beading, in turn, modulates the elevated [Ca^2+^]_axon_. We found that the stress-induced [Ca^2+^]_axon_ was highly heterogeneous. As resolved in the kymograph of the stressed axon (Fig. 5I, right), the [Ca^2+^]_axon_ inside the “beads” increased significantly. In contrast, inside the “strings”, it decreased significantly (Fig. 5J) after the flux onset, indicating such dramatic Ca^2+^ heterogeneity is driven by the sudden contraction of the axon walls in the stressed axons (Fig. 5K). After the flux, the Ca^2+^ trapped in the beads was released upon the relaxation of the flanking “string” regions (Fig. 5I, asterisk and Video 8). Notably, the transmission rates of the [Ca^2+^]_axon_ along the fluxed and beading axon was significantly reduced as compared to that along the non-beading axon prior to the flux (Fig. 5, L and M). These data demonstrate that mild mechanical stress induces the rapid contraction of the axon segments, which narrows the axon shaft to restrict the [Ca^2+^]_axon_ into the beading areas, impeding the long-range spreading of the [Ca^2+^]_axon_ beyond the mechanically stressed regions.

### Reversible axon beading is mechanoprotective against mild mechanical stress

To examine whether disruption of actomyosin-II affects the kinetics of [Ca^2+^]_axon_ in the stressed axons, we analyzed the [Ca^2+^]_axon_ kinetics following treatment with either NM-II inhibitor Para-Nitro-Blebbistatin (PNB), the low photo-toxicity analogue of Blebbistatin (Képiró et al., 2014), or NM-II inhibitor ML-7. We found that although both NM-II inhibitors abolished the formation of axonal beads that were induced by the low-speed flux (50 μL/min, 180s), in non-treated neurons, ML-7 disturbed the resting [Ca^2+^]_axon_ level (Fig. S5, D and E). We, therefore, chose to use PNB as the actomyosin-II inhibitor in the following experiments. While in control axons, the elevated [Ca^2+^]_axon_ was highly restricted to the beading regions (Fig. 6, A and B, left panels; and Video 9), the elevated [Ca^2+^]_axon_ in the PNB-treated axons was less restricted, showing a significantly wider distribution (Fig. 6, A and B, right panels; and Video 9). In PNB-treated neurons, the amplitude of the [Ca^2+^]_axon_ elevation along the entire axon region (Fig. 6, C and D) and the beading region (Fig. 6E) were both reduced compared to those of control axons (see also Video 9), suggesting that axonal beading is required for the [Ca^2+^]_axon_ elevation. Notably, in control axons, the [Ca^2+^]_axon_ elevations declined rapidly after the mild stress, as reflected by the sharp declining curve after the [Ca^2+^]_axon_ reached its peak (Fig. 6, E, black curve and F), while in the PNB-treated axons, the declining rate was significantly slower (Fig. 6, E, red curve and F), suggesting that actomyosin-II inactivated axons need more time to remove the elevated [Ca^2+^]_axon_ to its baseline after the stress. Besides the changes in the [Ca^2+^]_axon_ amplitudes, in PNB-treated axons, the occurrence of multiple [Ca^2+^]_axon_ bursts, which are sudden shooting of [Ca^2+^]_axon_ waves along the stressed axons bidirectionally (Fig. 6B, red arrowheads; Video 9, right panels) was significantly increased (Fig. 6G), indicating that the inactivation of actomyosin-II-dependent axon beading can significantly alter the kinetics of [Ca^2+^]_axon_ elevation, leading to the [Ca^2+^]_axon_ dysregulation, which is marked by the slower recovery rate and more frequent bursting rate of the elevated [Ca^2+^]_axon_.

**Figure 6.**
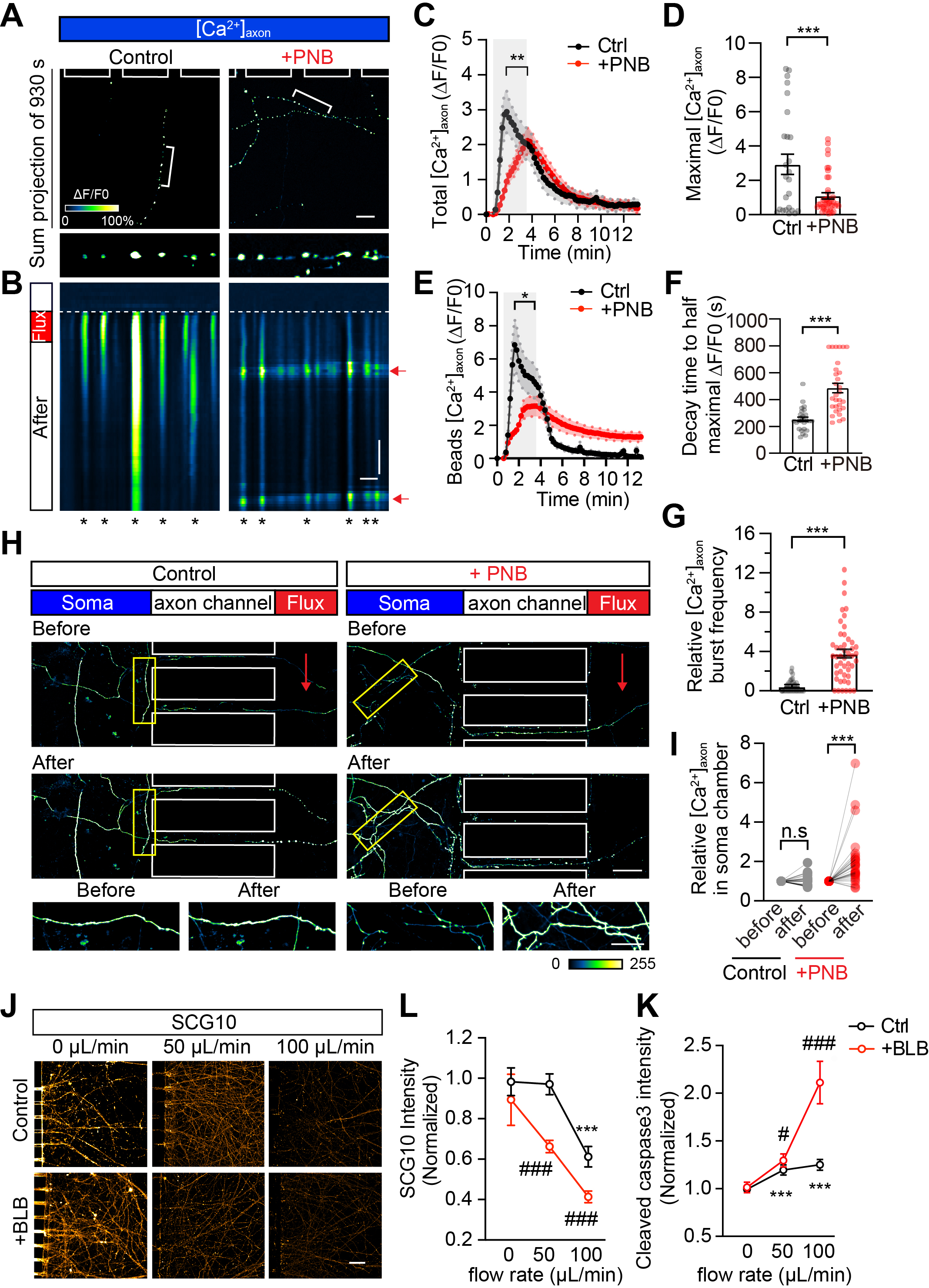
Actomyosin-II-dependent axon beading suppresses the spreading and bursting of the [Ca^2+^]_axon_ along the stressed axon. **(a)** [Ca^2+^]_axon_ hot spots before and after treatment with Para-Nitro-Blebbistatin (+PNB; 50 μμ). Bracketed regions are shown magnified in the lower boxes. The changes in fluorescence intensity (ΔF/F0) that reflect the kinetics of [Ca^2+^]_axon_ fluctuations are colour-coded. Bar = 20 μm. **(b)** Kymographs of magnified axons in **(a)**, the bursts of [Ca^2+^]_axon_ wave are indicated by red arrows on the right-hand side. [Ca^2+^]_axon_ hot spots are indicated with asterisks. x-axis Bar = 5 μm, y-axis Bar = 200 s. **(c-g)** Quantification of the ΔF/F0 changes reflecting the kinetics of [Ca^2+^]_axon_ fluctuations: **(c)** in the total axonal area (N = 26, 42, respectively), **(d)** the peak values of **(c)** (N = 26, 42, respectively)**, (e)** in the beading regions (N = 28, 30, respectively), **(f)** the time required to decay to half-maximal ΔF/F_0_ using the data set in **(e)** (N = 28, 30, respectively) and **(g)** the relative burst frequency of [Ca^2+^]_axon_ (N = 28, 47, respectively). **(h)** Representative tiled live-cell images showing the [Ca^2+^]_axon_ in the non-stressed (Soma) and stressed (Flux) regions. Yellow-boxed regions are magnified in the lower panels, with red arrows indicating the flux direction. [Ca^2+^]_axon_ intensity is colour-coded. Bar = 40 μm (top) and Bar = 20 μm (bottom). **(i)** Comparison of [Ca^2+^]_axon_ in the non-stressed soma chamber. N = 33, 30, respectively. **(j)** In control or BLB-treated neurons, the SCG10 intensity in the stressed axonal regions follows the stress induced by the indicated flow rates. Bar = 150 μm. **(k-l)** Quantification of SCG10 **(k)** and cleaved caspase 3 **(l)**. For SCG10 (control, N = 80, 105, 71, 63; +BLB, N = 90, 82, 84). For cleaved-Caspase 3 (control, N = 119, 131, 125, 105; +BLB, N = 84, 109, 114). Data represent mean ± s.e.m.; In (c-g, k, l) unpaired two-tailed student’s *t*-test; In (i) paired student’s *t*-test; \**p*<0.05, \*\**p*<0.01, \*\*\**p*<0.001; * marks the *t*-test results with the 0 μL/min control dataset; # marks the *t*-test results between control and BLB-treated neurons of the same flow rate.

As axon beading impedes long-range [Ca^2+^]_axon_ transmission, we further compared the intensities of [Ca^2+^]_axon_ in axons inside the soma chamber (Fig. 6H, yellow boxes), which were not stressed. We found that in PNB-treated groups, the [Ca^2+^]_axon_ intensity in the axons in the soma chamber was significantly increased following stress (Fig. 6I, red spots). In contrast, the control groups remained stable (Fig. 6I, grey spots). This result suggests that actomyosin-II inactivation increased the long-range spreading of elevated [Ca^2+^]_axon_ from stressed to non-stressed axonal regions (see also Video 9). The fate of these stressed axons in the AoC was also examined. We found that compared to the control axons stressed by the same rates of medium flux, the axons of Blebbistatin-treated neurons underwent degeneration to a degree that was dramatically more severe, represented by decreased SCG10 (Fig. 6, J and K, red curves) and increased cleaved Caspase 3 (Fig. 6L, red line). These data suggest that actomyosin-II is required to protect the mildly stressed axons from primary injury and subsequent degeneration.

### Up-regulating actomyosin-II activity alleviates axonal injury in mTBI mice

Finally, we investigated the function of stress-induced axon beading *in vivo* using a mouse model of mTBI. For this, we utilized closed-skull mouse models for TBI to mimic concussion, as previously described (Gu et al., 2017; Sun et al., 2022). Briefly, adeno-associated virus (AAV) encoding the vehicle, either the inactive MRLC mutant S19A/T18A (SA), the constitutively active MRLC mutant S19D/T18D (SD), or MRLC wild-type (WT) were injected into the primary somatosensory cortex of adult mice to manipulate the activity of actomyosin-II of the infected cortical neurons (Fig. 7A). Two weeks after the AAV injection, a single strike of impact was given on the secondary motor cortex in the opposite hemisphere of the AAV injection site (Fig. 7, B and C). After a 24-hour interval, the morphologies of the AAV-infected commissural axons in the tracts crossing the mid-line (corpus callosum) and those projecting to the site of impaction (impacted site) were analyzed (Fig. 7D). We found that in the impacted mice, more axons in both the corpus callosum and the impacted site demonstrated a fragmented appearance, marked by the shortened and sphere-like shapes (Fig. 7E, right panels) compared to sham mice (Fig. 7E, left panels). After comparing the morphological features of the stressed axons (Fig. 7F), we found that in both regions (corpus callosum and impacted site), the ratio of the fragmented axon was significantly increased in the impacted mice (Fig. 7G), showing that the mTBI model was successful. We next compared the severity of axon injury in mice brains with neurons expressing the MRLC mutants. We found that in mice brains infected with AAV expressing the constitutively active MRLC mutant S19D/T18D (SD), the proportion of fragmented axon was significantly reduced compared to the vehicle (-) or MRLC wild-type (WT) AAV-infected mice (Fig. 7, H and I), suggesting that activation of actomyosin-II inhibits axonopathy in the mTBI.

**Figure 7.**
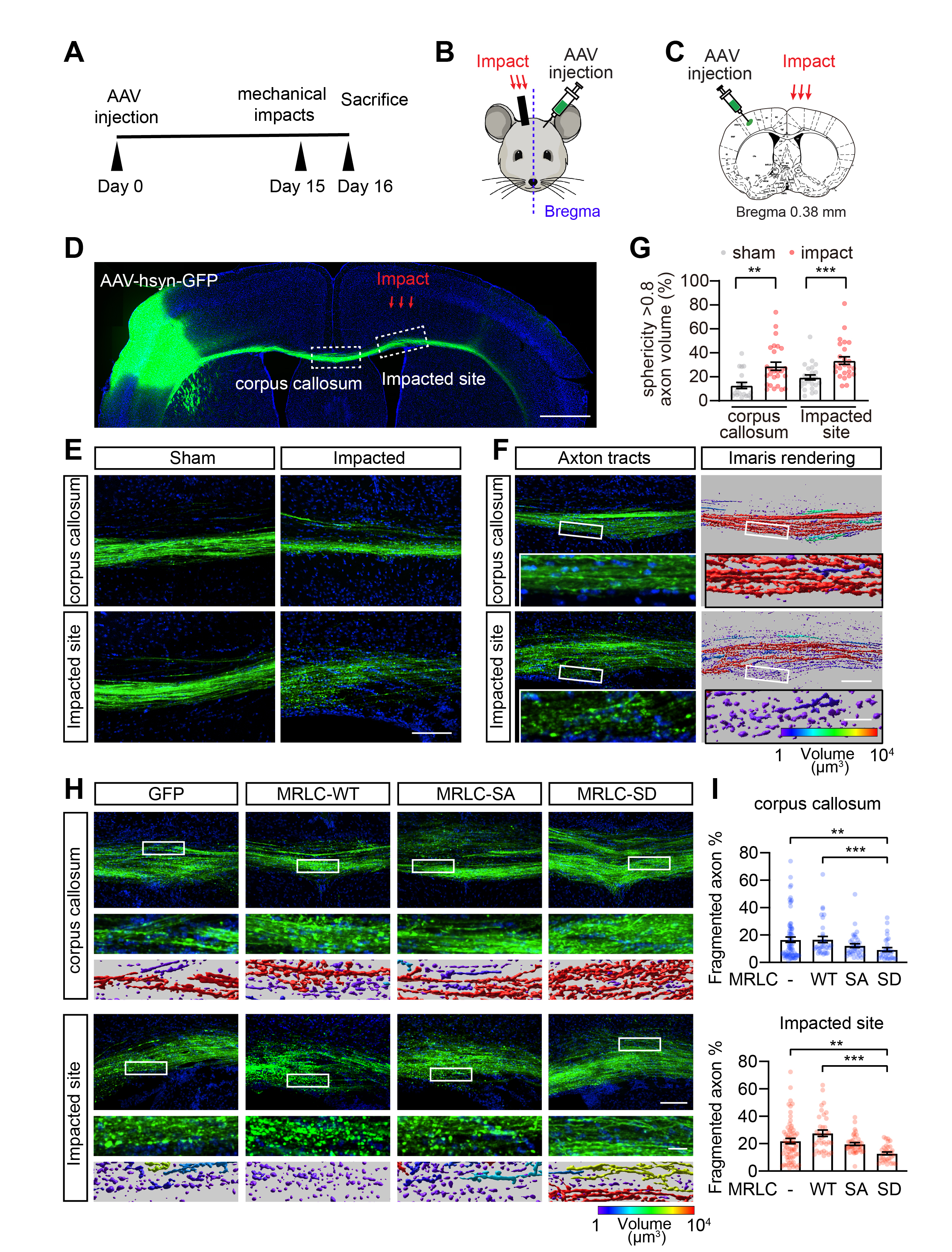
Actomyosin-II-dependent axonal beading protects mechanically stressed axons from degeneration. **(a)** Procedure timeline of the closed-skull mild Traumatic Brain Injury (mTBI) mouse model. **(b)** Schematic showing the AAV injection and impact sites in the brain of the mTBI mouse model. **(c)** Brain map showing AAV-hsyn-GFP injection and impact locations. **(d)** Representative tiled image showing axon tracts in the corpus callosum and impacted sites. Bar=2 mm. **(e)** High magnification images in the corpus callosum and impacted sites. Bar = 100 μm. **(f)** Z-stack raw images of axon tracts with 3D Imaris renderings of selected axonal segments shown on the right. White boxed outlines are magnified below. The volume of the axonal segments is colour-coded. Bar = 100 μm (top) and 20 μm (bottom). **(g)** Quantification of the fragmented axon (sphericity > 0.8; length < 10 μm) in **(f)**. N = 18, 26, 24, 25, respectively. **(h)** Representative images show the axons of neurons expressing either MRLC^WT^, MRLC^SA^, MRLC^SD^ or empty vector in commissural axons crossing the corpus callosum (top) or projecting to the impacted site (bottom) in the brain of the mTBI mice, the volume of axonal fragments is colour-coded. Bar = 100 μm (top) and 20 μm (bottom). **(i)** Percentage of fragmented axons (sphericity > 0.8; length < 10 μm) for the corpus callosum (top) and impacted site (bottom) in **(h)**. N = 73, 34, 37, 31, respectively (top); N = 70, 35, 40, 33, respectively (bottom). Data represent mean ± s.e.m.; unpaired two-tailed student’s *t*-test; \*\**p*<0.01, \*\*\**p*<0.001.

As shown in Fig. 8, by combining an AoC device, time-lapse super-resolution microscopy, and a mice TBI model, we discovered a new form of structural plasticity in CNS neurons that enables them to resist mild mechanical stress. The stress-induced reversible axon beading within the stressed region of the neuron is generated by actomyosin-II-driven radial contraction, which contracts the axonal cortex to prevent elevated [Ca^2+^]_axon_ from spreading beyond the stressed region to unstressed regions. Our loss- and gain-of-function experiments demonstrated that actomyosin-II in the axonal cortex plays a mechanoprotective role against mechanical stress in CNS neurons. Therefore, our findings suggest activating this machinery may help reduce primary axon injury in the CNS.

**Figure 8.**
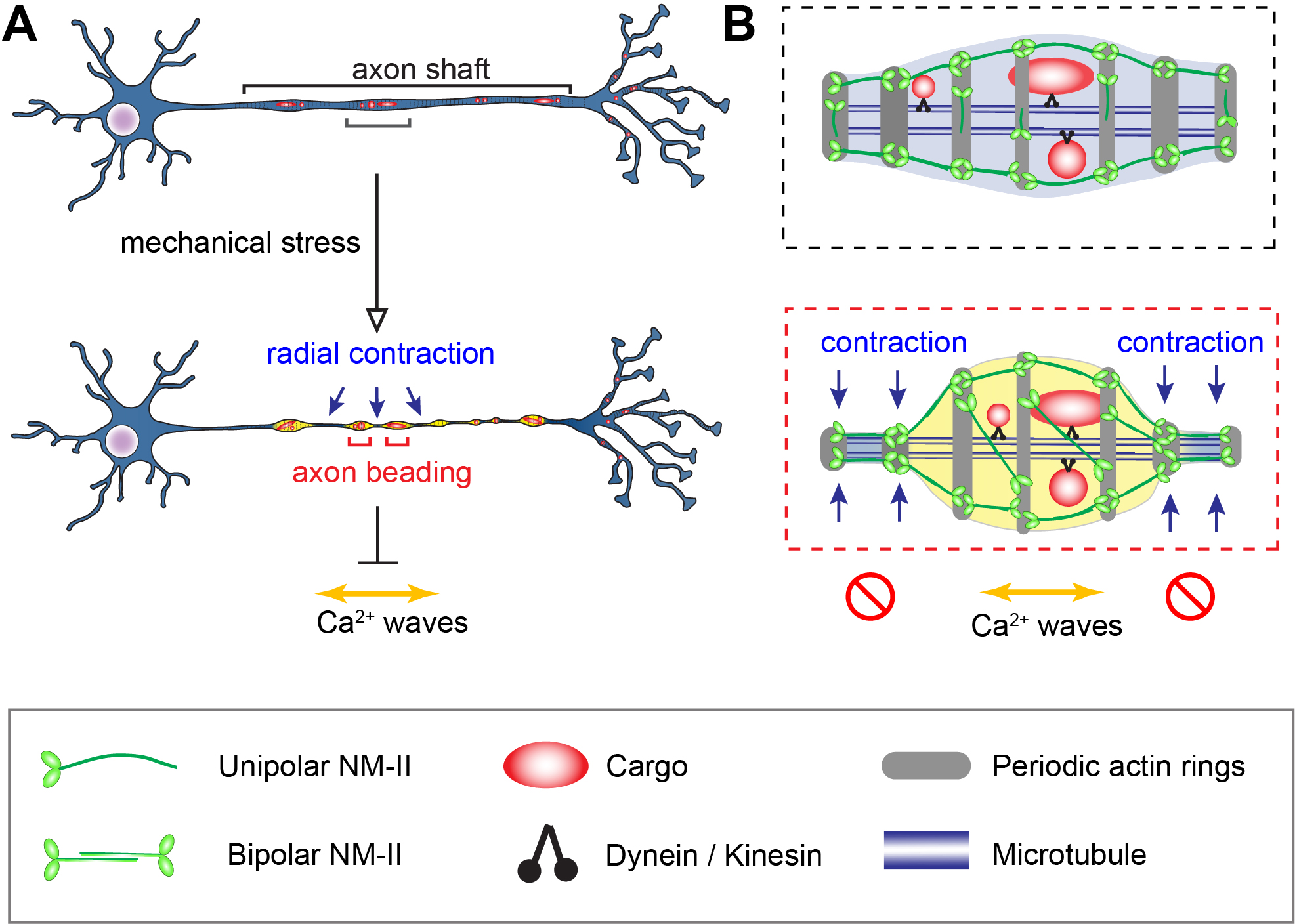
Actomyosin-II-dependent structural plasticity shields CNS axons from mild mechanical stress. **(a)** Actomyosin-II-driven radial contraction (blue arrows) leads to rapid and reversible axonal beading (red brackets) in axon shafts exposed to mild mechanical stress. The acute shape change confines the spreading of elevated Ca^2+^ (yellow regions within beads) in the axon, thereby protecting the stressed axon from widespread Ca^2+^ (yellow bidirectional arrow) and subsequent severe injury caused by mild mechanical stress. **(b)** Schematic of a single bead magnified to show how the acutely contracted axon restricts both organelle trafficking and Ca^2+^ spreading. Cellular components are indicated on the graph.

## Discussion

The extreme length of white matter axons dramatically increases their risk of being injured by mechanical stress *en route* (Cavanagh, 1984). Indeed, CNS axons are generally regarded as selectively vulnerable to mechanical impacts (Johnson et al., 2013; Smith and Meaney, 2000). However, it remains unknown why CNS axons can resist a significant extent of surface compression during daily activities and contact sports, which causes significant deformation of the soft brain (Funk et al., 2011; Knutsen et al., 2020). Our study outlined a new proactive function of the periodic axonal actomyosin network in shielding axons from mild transverse forces *in vitro* and *in vivo*.

### Radial contraction of the axon cortex promotes their resistance to mild mechanical stress

Our AoC model combined with super-resolution microscopy has allowed us to separate the primary axonal responses from the secondary cascades of events leading to degeneration, such as those that occur in distal axons during Wallerian degeneration (Ding and Hammarlund, 2019a). The use of time-lapse super-resolution microscopy allowed us to precisely delineate the subcellular responses to discrete levels of forces transversally applied to axons. Our results reveal that actomyosin-II mediates a novel mechanoprotective response to mild transverse stresses (lower than 0.467 Pa). These perturbations consistently induce reversible axon beading driven by actomyosin-II-dependent acute radial contraction, dramatically and non-homogenously reducing axon diameter. This contractile effect stalls the axon organelle trafficking and suppresses the firing and long-range spreading of stress-induced Ca^2+^ waves. Recent simulation studies theoretically predicted a role for the axon cortical actin network in shielding deeper cellular structures from primary injuries caused by mechanical stimuli (Dubey et al., 2020; Gu et al., 2017; Kant et al., 2021; Sun et al., 2022). Our data provide solid evidence for this theory by showing that actomyosin-II-dependent contraction underlies the resistance against mild mechanical insults. Importantly, we provide an *in vivo* validation of the mechano-protective properties of the axonal actomyosin network with our mTBI mouse model, demonstrating that activation of actomyosin-II alleviates primary axon injury in the mouse brain. The newly identified mechano-protective function of actomyosin-II is highly in line with our findings that periodic actomyosin-II plays a critical role in maintaining the structural stability of CNS axons (Wang et al., 2020).

Therefore, the periodically distributed actomyosin-II drives the contraction of the axon cortex, acting similarly to a thin layer of muscle in shielding vulnerable axons from mechanical stresses. Targeting this intrinsic mechanoprotective machinery may be a new strategy for preventing mechanical axonal injury at the earliest stage of primary mechanical injury, providing more insightful knowledge for treating or mitigating mTBI and concussion.

### The stress-induced reversible axon beadings are not focal axonal swellings (FAS)

We have revealed that the reversible axon beading process differs from the FAS observed in the Wallerian degenerations (Ding and Hammarlund, 2019b). Our data indicate that the transient axon beads represent a plastic response of the axon evolved to resist mild mechanical stress. In contrast, FAS marks the permanent pathological damages detected in degenerating axons (Tang-Schomer et al., 2012; Valiyaveettil et al., 2014). They differ in three main aspects. Firstly, the stress-induced axonal beads are formed by contraction rather than passive swellings, reversible upon the relaxation of actomyosin-II. As shown in Fig. 2, the dynamic axonal beading is completed within the first 5 s following the onset of mild stress (less than 0.467 Pa) and is restored within 5-6 min after the withdrawal of the flux. Secondly, the MT tracks, axon surface and organelle trafficking machinery remain intact in the transiently appeared beads, as revealed by TEM, SEM and live-cell imaging microscope. These data demonstrate that organelles are paused by the radially contracted axon but are soon released upon relaxation of the cortical actin network to continue their trafficking. Thirdly, in axons bearing stress-induced beads, the long-range spreading of the elevated Ca^2+^ to the intact neuronal parts is hindered, leading to the regional restriction of injury signals. In contrast, in FAS, the Ca^2+^ surges induced by injury progressively propagate throughout the whole neuron, ultimately leading to degeneration.

Our study provides valuable insights into the modulation of stress-induced Ca^2+^ elevation in CNS axons. However, the exact source of the Ca^2+^ remains unclear. Previous studies have proposed that TRPV4 (Gu et al., 2017) and other non-specific Ca^2+^ channels could mediate the stress-induced Ca^2+^ currents (Gaub et al., 2020; Pan et al., 2022; Vargas et al., 2015; Witte et al., 2019). However, further efforts are needed to determine the source of the Ca^2+^ and examine whether targeting the downstream of the actomyosin-II could also be protective against mechanical stress. Additionally, several minutes after axon beading, we noticed a rapid decline in the elevated Ca^2+^. The mechanisms underlying such Ca^2+^ clearance also remain unknown. Future investigations into the mechanism allowing rapid Ca^2+^ clearance are needed. Finally, whether extracellular cues can activate actomyosin-II-dependent mechanoprotective machinery to improve axonal resilience remains untested. Additional screening approaches are warranted to elucidate upstream signalling pathways that activate actomyosin-II and explore its potential therapeutic applications.

## Methods

### Reagents, Antibodies and DNA constructs

Blebbistatin (#ab120425, Abcam); Para-Nitro-Blebbistatin (#24171, Cayman); ML-7 (#11801, Cayman); Calyculin A (#9902S, Cell Signaling); BAPTA-AM (#T6245, Targetmol); Nocodazole (#M1404, Merck); Taxol (#S1150, Selleck); EDTA (#A500895, Sangon Biotech). The following primary and secondary antibodies were purchased from commercial suppliers: Anti-Cleaved Caspase 3 (Cell Signaling, #9661S); anti-SCG10 (STMN2) (Proteintech, #10586-1-AP); anti-β III tubulin (Proteintech, #66240-1-Ig-100ul); anti-β III tubulin (merck, #AB9354); anti-Phospho-Myosin Light Chain 2 (Thr18/Ser19) (Cell Signaling, #3674); anti-Myosin Light Chain 2 (Cell Signaling, #8505); anti-βII-Spectrin (SantaCruz, #sc-136074); anti-GAPDH (Proteintech, #10494-1-AP); anti-GFP (Aves Labs, #GFP-1020); anti-NM-IIB αCT (Sigma-Aldrich, #M7939); anti-NM-IIB αNT (Santa Cruz, #sc-376954); Alexa Fluor 488 donkey anti-chicken IgY (IgG) (H+L) (Jackson, #703-545-155); DyLight 488 Goat anti-mouse IgG (Thermo Fisher Scientific, #P36961); Alexa Fluor 568 goat anti-rabbit IgG (Thermo Fisher Scientific, #A11011); Alexa Fluor 568 goat anti-mouse IgG (Thermo Fisher Scientific, #A11031); Alexa Fluor 568 goat anti-Chicken IgY (Abcam, #ab175477); Alexa Fluor 647 goat anti-mouse IgG (Thermo Fisher Scientific, #A32728); Alexa Fluor 647 goat anti-rabbit IgG (Thermo Fisher Scientific, #A21244); HRP-labeled Goat Anti-Rabbit IgG (Abcam, #ab131366); HRP-labeled Goat Anti-Mouse IgG (Abcam, #ab131368). The primers GFP-forward (CGAAGGCTACGTCCAGGAGC) and GFP-reverse (CGATGTTGTGGCGGATCTTG) were synthesized from Tsingke, Shanghai, for the AAV virus titration. The DNA construct encoding Lifeact-G/RFP was provided by Roland Wedlich Soldner (MPI Biochemistry, Martinsried); pTagRFP-mito was purchased from Evrogen (#FP147); GCaMP-6f, pEGFP-MRLC (Plasmid #35680), pEGFP-MRLC T18AS19A (Plasmid #35681) and pEGFP-MRLC T18DS19D (Plasmid #35682) were purchased from addgene; pAAV-hsyn-GFP was a gift from Professor Zhenge Luo (Shanghaitech University, Shanghai).

### Neuronal culture and transfection in the AoC

The polydimethylsiloxane (PDMS) slab of the AoC was fabricated as described in a previous study (Pan et al., 2022). Then the slab was plasma-treated, sealed against a glass-bottom dish, and immediately immersed in ethanol to enhance the hydrophilicity of the PDMS surface. Under sterile conditions, a glass-bottom dish attached to the PDMS microfluidic device was washed with 100% ethanol, then Phosphate buffered saline (PBS) 3 times before being coated with 0.5 mg/mL Poly-L-Lysine in Borate buffer (1M, pH 8.5) in all chambers overnight. Hippocampal neurons were cultured from embryonic day 18 (E18) embryos of Sprague Dawley rats. All experiments were performed following relevant guidelines and regulations as approved by the Animal Ethics Committees of ShanghaiTech University (approval number: 20230217002) and the Chinese Academy of Sciences (approval number: NA-058-2021). Hippocampal neurons were prepared and seeded into the soma chambers of microfluidic devices at a minimum of 2×10^5^ cells per reservoir. Lipofectamine 2000 was used to transfect the neurons with indicated plasmids on DIV5 to 6. Flux-induced injury and stress experiments in the AoC were performed on DIV 7 to 8.

### AAV package and mTBI mice model

The plasmids MRLC mutant S19AT18A (SA), MRLC mutant S19DT18D (SD), and MRLC wild-type (WT) were modified with a P2A sequence at their N terminus and then cloned into the pAAV-hsyn-GFP vector. AAV viruses were produced by co-transfecting HEK293T cells with the AAV helper plasmid, PHP.s (Chan et al., 2017), and the transgene plasmids mentioned before using the PEI transfection reagent. The transfected cells were incubated for 72 hours for AAV production, and the cell pellet was resuspended in lysis buffer (150 mM NaCl, 20 Mm tris, pH8.0). After being treated with Benzonase for 15 min, the supernatant containing AAV particles was collected by centrifugation. AAV particles were purified using OptiPrep-density gradient ultracentrifugation (stemcell, 07820) in 40% fraction. The titer was determined using qPCR with primers targeting the GFP region. The qPCR primers for GFP are listed as follows: forward, 5′-CGAAGGCTACGTCCAGGAGC-3′; reverse, 5′-CGATGTTGTGGCGGATCTTG-3′. The purified AAV particles were used for *in vitro* and *in vivo* experiments. AAV virus injection was slowly injected into the primary somatosensory cortex (M/L +3.0/3.3/3.5 mm; A/P +1/+0.5/-0.5mm; D/V-0.6/-0.4mm) in 8-week-old wild type C57BL/6J mice. The mild TBI experiments were performed two weeks after the injection. The C57BL/6J mice (N=16) were put into a stereotaxic frame with rounded head holders after being given 5% isoflurane anesthesia. Isoflurane was delivered by nose cone at 2% in air. After shaving the heads, a midline skin incision was made to reveal the skull. An electromagnetic stereotaxic impact device with a rubber tip (3 mm in diameter) was used to impact the secondary motor cortex (M/L −0.5 to −3.0 mm; A/P +2 to −1mm; D/V −2.4mm) at a speed of 3.5 m/s for a duration of 100 ms. The mice were immediately sacrificed, perfused and fixed after 24 hours.

### Brain slice sectioning, IF staining, imaging and analysis

Mice brains were subjected to cardiac perfusion and fixation. The fixed brains were left overnight in 4% paraformaldehyde and subsequently placed in 30% sucrose for at least 24 hours for dehydration. The dehydrated brain tissue was then submerged in OCT (SAKURA, #4583) in a plastic mould, avoiding air bubbles, and frozen quickly in liquid nitrogen. The frozen tissue was then sectioned into 40 µm slices using a Leica cryostat microtome CM3050S and collected onto slides. For Immunofluorescence, the samples were blocked with 5% donkey serum in PBST for 1 hour, followed by adding primary antibody GFP (1:1000; Aves Labs, #GFP-1020), which was incubated overnight at 4°C. The slides were then incubated with donkey anti-chicken secondary antibody Alexa Fluor 488 (1:5000; Jackson, #703-545-155) for 2 hours at room temperature. Samples were immediately mounted in fluoroshield mounting medium (Sigma-Aldrich, #F6182) for imaging. The brain slices were imaged using the Nikon inverted spinning confocal microscope (CSU-W1) with a 20×0.75 NA objective. Using up to 40 cycles of iterative deconvolution, 3D stacks of confocal images were deconvoluted in the Huygens Professional (v18.10, Scientific Volume Imaging). Then, Imaris (Imaris 9.7.2, Bitplane) was used to render the fluorescent axonal segments into “surface” detected in manual mode with the following parameters: surface grain size= 0.800 μm, diameter of largest sphere= 2.800 μm, manual threshold value= 2.969. After the rendering, the fragmented axon segments were extracted by setting the filter of the sphericity to more than 0.8 and the length to less than 10 μm.

### Live-imaging and automatic analysis of axons in AoC

We conducted high-temporal-resolution imaging of axons experiencing medium flux in the AoC using a Nikon TI2-E inverted microscope with a Yokogawa spinning confocal disk head (CSU-W1) and a 63×1.4 NA/219.15 µm WD/0.1826 µm/pixel (1,200 × 1,200) objective. To assess the effects of various compounds on axon beading, we prepared Blebbistatin (50 μμ), Para-Nitro-Blebbistatin (50 μμ), ML-7 (10 μμ), Calyculin A (50 nM), latrunculin B (5 μμ), BAPTA-AM (10 μμ), Nocodazole (50 μμ) and Taxol (10 μμ) in DMSO, and EDTA (0.5 mM) in sterilized Milli-Q water. These compounds were added to the neuron culture medium of the AoC 30 minutes before the microfluidic flux session. We used Neurobasal minus phenol red (#12348017; Thermo Fisher Scientific) as the imaging medium. CellMask DeepRed (Invitrogen) was added to the culture medium at a final concentration of 5 µg/mL two hours before the flux session. Variable microfluidic flux was generated by injecting conditioned culture medium into the central injury channel using Pump 11 Elite Programmable Syringe Pumps (Harvard Apparatus, #704505) at indicated flow rates for 180 seconds while time-lapse images were acquired simultaneously. The axon beading in time-lapse images was automatically analyzed using ImageJ software (National Institutes of Health, v2.0.0/1.52p). The [Ca^2+^]_axon_ intensity analysis in time-lapse images was performed using ImageJ software (National Institutes of Health, v2.0.0/1.52p). Firstly, background noise was removed using the “subtract background” plugin with the rolling ball radius set to 200 pixels. Then, axons were outlined by the “segmented line” tool. Next, the [Ca^2+^]_axon_ intensity values along the line were measured using the “line profile” plugin. Kymographs were generated using the Multi-Kymograph plugin for ImageJ. All measurements were analyzed using GraphPad Prism. Finally, all images and figures were compiled using Illustrator CS 23.1.1 (Adobe).

### Time-lapse Lattice SIM and analysis

The F-actin marker plasmid Lifeact-GFP was transfected into the neurons cultured in AoC, and on DIV8, the cells were subjected to medium flux (50 μL/min) for 180s while being live-imaged using the Structured Illumination Microscope (ZEISS Elyra 7 with Lattice SIM). The imaging was performed using the 63×1.4 NA oil immersion objective with 15 frame accumulations as multi-slice z-stacks during the time-lapse set at 90s intervals for 16 minutes. The 3D time-lapse raw images were processed using the adjusted mode of the SIM algorithm in Zen software, which included a strong sharpness filter and the fast fit advanced filter. Next, the Z-stack microscopic images of live neurons were exported to Imaris software (version 9.7.2; Bitplane) for analysis. The extent of radial contraction was analyzed using the ‘filament’ function. Axon filaments were manually rendered by fitting the axon diameter with a lower contrast threshold of 4.500 and using the “Approximate Circle of Cross Section Areas” mode. The ‘Filament Analysis’ plugin was used to cut the filaments into continuous spots with diameters corresponding to the fluctuated diameter along the axon. The spots were colour-coded from 0.15 µm to 2.10 µm. The changes in the diameter distribution of spots constituting the axon were quantified to determine the extent of radial contraction and dilation. For 3D-SIM images, the same process was followed, but the axon diameter was fitted with a lower contrast threshold of 0.08, and the continuous spots were colour-coded from 0.10 µm to 2.50 µm

### STED Microscope and data processing

On DIV9, hippocampal neurons were treated with Blebbistatin (BLB, 50 µM) or Calyculin A (CA, 50 nM) for 30 minutes before being fixed and imaged using a Leica TCS SP8 STED 3X Microscope equipped with a tunable white light laser and 775nm, 660nm, 592nm STED depletion lasers. To prepare the samples for 3D-STED imaging, they were initially fixed in 4% paraformaldehyde for 30 minutes at room temperature and then blocked with a blocking buffer (0.1% saponin, 1% BSA, 0.2% gelatin in PBS) for 1 hour. The primary antibody NM-IIB αCT (1:1000) and NM-IIB αNT (1:500) were added to the dish and incubated at 4°C overnight. Goat anti-rabbit secondary antibody Alexa Fluor 647 (1:5000) and Goat anti-mouse-488 secondary antibody DyLight 488 (1:500) were incubated for 1 hour at room temperature. For Dual-colour STED imaging, samples were immediately mounted in ProLong Diamond antifade medium (Thermo Fisher Scientific, P36961). Images were acquired using the HC plan apochromat 100x 1.4 NA oil immersion objective using Leica LASX software with 11 frame accumulations as multi-slice z-stacks. NM-IIB αNT was excited at 495 nm, STED depleted at 592 nm with the laser power of 40%, while the NM-IIB αCT was excited at 631 nm, and STED depleted at 775 nm with the same laser power. The resulting 3D stacks of STED images were deconvoluted and z-drift corrected in Huygens Professional (Version 18.10, Scientific Volume Imaging, the Netherlands) using up to 40 cycles of iterative deconvolution. The fluorescent volumes in the deconvolved Z-stack images were rendered into “spots” using Imaris software (Imaris 9.7.2, Bitplane). After subtracting the background, spots were manually detected using the following parameters: estimated XY diameter = 0.120 µm, estimated Z diameter = 0.400 µm. NM-IIB filaments were extracted by setting the filter of the distance between the centre of the spots to less than 0.120 µm using the Imaris plugin “Colocalize Spots”. The unipolar or bipolar NM-II filaments were further filtered based on the number of connected spots.

### SEM and TEM

The DIV8 rat hippocampal neurons in AoC were fluxed at a flow rate of 50 μL/min for 180 s. Following the flux, all neurons were fixed in 2.5% glutaraldehyde for 2 hours at 4℃. The coverslips were removed and transferred to 1% osmium tetroxide for 2 hours at 4℃. To prepare for SEM, the samples were dehydrated using an ascending ethanol series (30%-100%) at 4℃ and then critically dried using a low-speed critical point dryer (Leica EM CPD300). Subsequently, the samples were attached to SEM pin studs with conductive double-sided carbon tape and coated with 8 nm gold layers using a vacuum sputter coater (Leica EM ACE200 Low Vacuum Coater). The gold-coated samples were imaged using a Zeiss field emission scanning electron microscope (GeminiSEM 460) equipped with a Gemini 2 electron optical column and SE2 detector, with an accelerating voltage of 2.0 kV and a beam current of 50 pA. For TEM, after secondary fixation with 1% osmium tetroxide for 2 hours at 4℃, the cells were stained with 1% uranyl acetate overnight at 4℃. The samples were dehydrated with ethanol and gradually infiltrated with liquid resin. The resin was polymerized, and sections (50 nm) were cut using an Ultramicrotome Leica EM UC7. The sections were observed using a GeminiSEM 460 microscope with a STEM detector, operated at 15.0 kV accelerating voltage and 1.0 nA.

### Statistical information

We used GraphPad Prism (GraphPad Prism v9.3.1) for statistical analyses. All measurements were taken from distinct samples. Results are reported as mean ± s.e.m. Outliers were removed using the ROUT method at Q=1% in Prism. For group comparisons, two-tailed nonparametric *t*-tests were executed. P values < 0.05 indicated statistical significance. Pilot experiments were performed to determine the requirements for sample size. Sample size sufficiency was determined by preliminary data or discussion. The sample size was always independently performed three or more times for statistical significance. Anderson-Darling test (alpha = 0.05) was used to test the normality of data sets. Locations of ROI for imaging acquisition were randomly selected in all experimental groups. Unless specified, all experiments were subjected to three biological replicates and two technique replicates, and all attempts at replication were successful. Data collection and analysis were performed by different operators, who were blind to the conditions of the experiments.

## Supplemental material summary

**sFig. 1** shows that the flow rate did not affect the formation and recovery rate of the axon beading process and supports Fig. 1. **sFig. 2** reveals that the radial contraction induced by the flux alters the morphology of axonal organelles and supports Fig. 2. **sFig. 3** shows that the microtubule tracks do not contribute to the reversible axon beading process and supports Fig. 3. **sFig. 4** shows that the flux-induced axon beading and Ca^2+^ elevation were regionally restricted in the stressed axonal segments, supplementing Fig. 5. **sFig. 5** shows that the resting Ca^2+^ level in axons is affected by the treatment of actomyosin-II inhibitor ML-7, supporting Fig. 6. **Videos 1 and 2** show the time-lapse images of reversible axon beading and its automatic detection. **Video 3** shows the time-lapse SIM images revealing the diameter contraction of individual actomyosin rings induced by flux. **Videos 4 and 5** display the deformation of mitochondria and paused axon trafficking during the flux-induced axonal radial contraction. **Video 6** shows the flux-induced periodic ring contraction during the beading process. **Video 7** demonstrates the process of beading formation in axons treated with actomyosin-II inhibitor/activator during flux. **Videos 8 and 9** exhibits the dynamic Ca^2+^ wave transmission and stress-induced axon beading process during the flux.

## Data availability statement

All data reported in this paper will be shared by the lead contact upon request. This paper does not report the original code. Any additional information required to reanalyze the data reported in this paper is available from the lead contact upon request.

## Acknowledgements

We thank Prof. Zhenge Luo, Dr Xiaoming Li, Dr Ziwei Yang, Dr Xiuqing Fu, Chengyu Fan, Rui Wang, Dr Alex McCann and Dr Xu Wang for her/his expert technical assistance. This work was supported by the National Natural Science Foundation of China (31871036 and 32271001 to T. Wang) Y. Chu acknowledges the support from the National Natural Science Foundation of China (32100777). Y. Liu would like to thank the Double First-Class Initiative Fund of ShanghaiTech University (SYLPOC0022022, SYLDX0302022). F.A. Meunier acknowledges the support from NHMRC Senior Research Fellowship (GNT1155794), grant (GNT1120381) and the Australian Research Council equipment grant (LE130100078). The authors declare no competing financial interests.

## Author contributions

Conceptualization: T. Wang, Y. Li, CX. Zhao, Y. Liu; Methodology: X. Pan, J. Li, G. Lei, W. Zhan, T. Luan, Y. Hu; Investigation: X. Pan, J. Li, Y. Chu, Y. Feng; Visualization: X. Pan, Y. Hu, T. Luan; Tools: FA Meunier; Supervision: T. Wang, Y. Li, Y. Liu; Writing—original draft: T. Wang, X. Pan, T. Luan, Y. Hu; Writing—review & editing: Y. Li, CX. Zhao, Y. Liu, FA Meunier.

## Declaration of interests

The authors declare that they have no competing interests.

## Supplementary information

**Video 1. Reversible axon beading is caused by the low-speed flux in an AoC device.**

Rat hippocampal neurons cultured in an AoC device were transfected with Lifeact-GFP and subjected to mild mechanical stress by injecting low-speed flow (50 µL/min, 180 s) into the central injury chamber. Instant axonal deformation during flux-induced stress was monitored using time-lapse confocal microscopy. A representative movie shows the reversible axonal beading process, with the duration of flux indicated with a white arrow and cyan background in the top panel. The boxed region of interest (ROI) is shown magnified in the bottom panel. Bar = 20 µm (top), 5 µm (bottom left), and 3 µm (bottom right).

**Video 2. Automatic detection of the mechanical stress-induced axon beading in 3D z-stack images using Imaris.**

Rat hippocampal neurons cultured in an AoC device were transfected with Lifeact-GFP. The mild mechanical stress was induced by injecting low-speed flow (50 µL/min, 180 s) into the central injury chamber. Z-stack confocal images were acquired in the middle injury chamber before and after the flux. The ‘Filament’ function of Imaris software was used to render the 3D z-stack images into filaments, which were then divided into continuous spots using the plugin ‘Filament Analysis’. The final renderings are displayed as multiple continuous spots with diameter fluctuation colour-coded. Bars = 30 µm (left) and 2 µm (right; magnification).

**Video 3. Mild mechanical stress causes reversible axonal diameter contraction.**

In an AoC device, rat hippocampal neurons expressing F-actin marker Lifeact-GFP were stressed with 50 μL/min flux for 180 s. 3D-SIM time-lapse images were acquired in the central injury chamber showing the dynamic axonal diameter change. The axons were rendered into continuous spots using the plugin ‘Filament Analysis’ of Imaris software and displayed with diameter fluctuation colour-coded. The appearance of the cyan boxes and arrows marks the duration of the flux. Bars = 10 µm and 5 µm (magnified panel).

**Video 4. Mild mechanical stress causes the deformation of axonal mitochondria.**

In the AoC device, rat hippocampal neurons co-expressing Lifeact-GFP and TagRFP-mito were subjected to flux-induced mechanical stress (20 μL/min, 180 s). Dual-colour time-lapse images were acquired using a spinning disc microscope during the flux, showing the dynamic morphological changes of both the beading axon and the mitochondria. The appearance of cyan boxes marks the duration of the flux. The boxed ROI is amplified in the right panels, with the compressed mitochondria (magenta) within the axonal beads (green) indicated by arrowheads. Bars = 20 µm (left) and 3 µm (right).

**Video 5. Reversible axon beading affects the axonal trafficking of mobile mitochondria.**

Rat hippocampal neurons cultured in an AoC device were transfected with Lifeact-GFP and TagRFP-mito. Mild mechanical stress was induced by injecting low-speed flow (20 µL/min, 180 s) into the central injury chamber. Dual-colour time-lapse images were acquired using a spinning disc microscope during the flux, showing the dynamic morphological changes of both the beading axon and the mitochondria. The appearance of cyan boxes marks the duration of the flux. The boxed ROI is shown magnified in the right panels, with the mobile mitochondria (magenta) indicated by arrowheads. Bars = 10 µm (left) and 5 µm (right).

**Video 6. Mechanical stress affects the periodic actin ring diameter.**

In an AoC device, axons of rat hippocampal neurons expressing F-actin marker Lifeact-GFP were subjected to mechanical stress (50 μL/min, 180 s), which was injected into the central injury chamber. 3D-SIM time-lapse images show dynamic morphological changes of the periodic actin ring during the flux. The appearance of the cyan boxes marks the duration of the flux. The boxed ROI is shown magnified in the right panels, with the detailed actin ring diameter change indicated by arrowheads. Bars = 10 µm (left), 3 µm (top right) and 1 µm (bottom right).

**Video 7. Stress-induced axon beading requires actomyosin-II activity.**

Rat hippocampus neurons expressing Lifeact-GFP or Lifeact-RFP were pre-incubated with either Blebbistatin (50 μM), ML-7 (10 μM), Calycullin A (50 nM), or latrunculin B (5 μM) for 30 min. The axons of these neurons were then subjected to flux-induced mechanical stress (50 μL/min, 180 s) in an AoC device. Time-lapse images were acquired and processed automatically using ImageJ Macro (see also Methods), which separated and quantified the “beading” (magenta) and the non-beading “between” (cyan) regions. Bar = 10 µm.

**Video 8. Stress-induced axon beading restricts the spreading of elevated Ca^2+^ in the axon.**

Rat hippocampal neurons expressing GCaMP-6f were stressed with 50 μL/min flux for 180 s in an AoC device. Time-lapse images were acquired, showing the elevated [Ca^2+^]_axon_ induced by flow injection. The appearance of cyan boxes marks the duration of the flux. The elevated Ca^2+^ restricted within a single axonal bead is marked by an asterisk. The spreading of the Ca^2+^ signal after the relaxation of the contracted regions flanking this axonal bead is indicated by arrowheads. Bars = 10 µm (left) and 5 µm (right).

**Video 9. Actomyosin-II-dependent radial contraction of axons is required to restrict stress-induced Ca^2+^ elevation.**

Axons of rat hippocampal neurons expressing GCaMP-6f were subjected to mild mechanical stress (50 μL/min, 180 s) in an AoC device. Time-lapse images of [Ca^2+^]_axon_ were acquired in the central injury chamber, showing the dynamic kinetics of [Ca^2+^]_axon_ in both controls (left) and PNB-treated (right) axons. The cyan background indicates the duration of the flux, and the boxed ROI is magnified in the lower panels, Bars = 20 µm (top) and 5 µm (bottom).

---

**Supplementary Figure 1.**
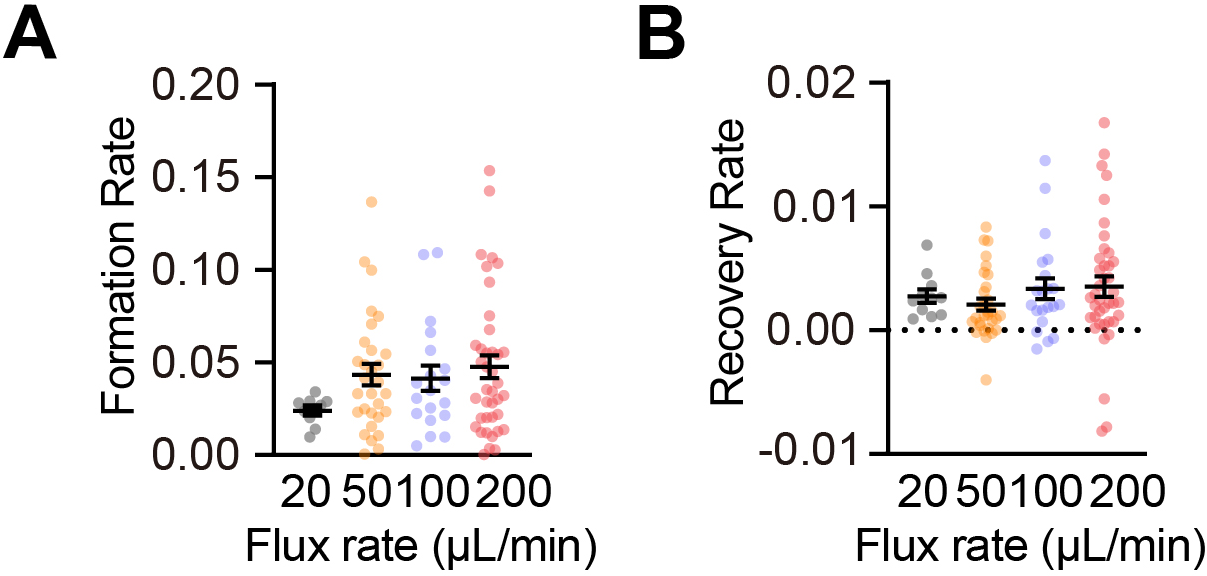
The degree of axon beading induced by flux injected into the AoC device varies with the flow rate. **(a)** Quantification of the formation rate of axon beading shown in Fig. 1e. N = 10, 30, 19, 39, respectively. **(b)** Quantification of the recovery rate of axon beading in Fig. 1e. N = 11, 30, 21, 40, respectively. Data represent mean ± s.e.m; unpaired two-tailed student’s *t*-test; n.s. Non-significant.

**Supplementary Figure 2.**
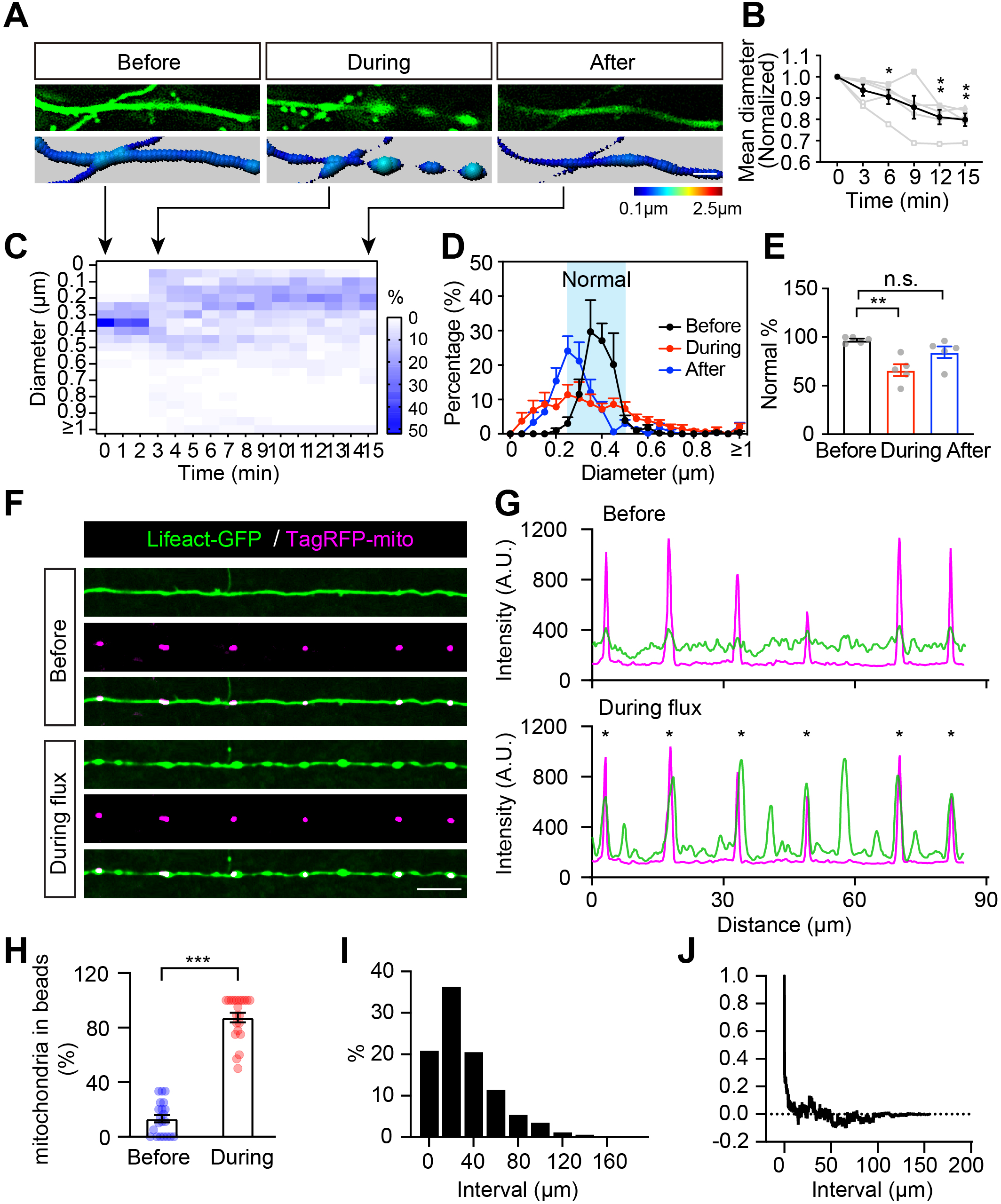
Flux-induced radial contraction of axons affects the trafficking and morphology of axonal organelles. DIV5-6 rat hippocampal neurons cultured in an AoC device were either transfected with Lifeact-GFP or co-transfected with both Lifeact-GFP and TagRFP-mito and live-imaged on DIV7-8 using a spinning-disc inverted confocal microscope. **(a)** 3D-SIM images were rendered into filaments using the Imaris software and displayed as multiple continuous spots with diameter fluctuation colour-coded. Bar = 2 µm. **(b)** The mean axon diameter of **(a)** is shown. N = 5. **(c)** Heatmap showing the frequency distribution of the diameter of an axon during the beading process. **(d)** The frequency distribution of axonal diameter before, during, and after flux is shown. **(e)** Quantification of normal axon segments with a diameter between 0.25 and 0.5 μm before, during, and after flux. N = 5. **(f)** The localization of the mitochondria is compared with that of the “beading” areas. Bar = 10 μm. **(g)** Line profile showing the fluorescent intensity of mitochondria (TagRFP-mito), and the axonal volume (Lifeact-GFP). Asterisks indicate the beads containing mitochondria. **(h)** Quantification of **(g)**, N = 21. **(i)** Distribution of the spacing intervals between beads. **(j)** The cross-correlation between these intervals. The axonal intervals were measured from 855 beads obtained from 42 axons. Data represent mean ± s.e.m.; In (b, e, h) paired two-tailed student’s *t*-test; \**p*<0.05, \*\**p*<0.01, \*\*\**p*<0.001; n.s. Non-significant.

**Supplementary Figure 3.**
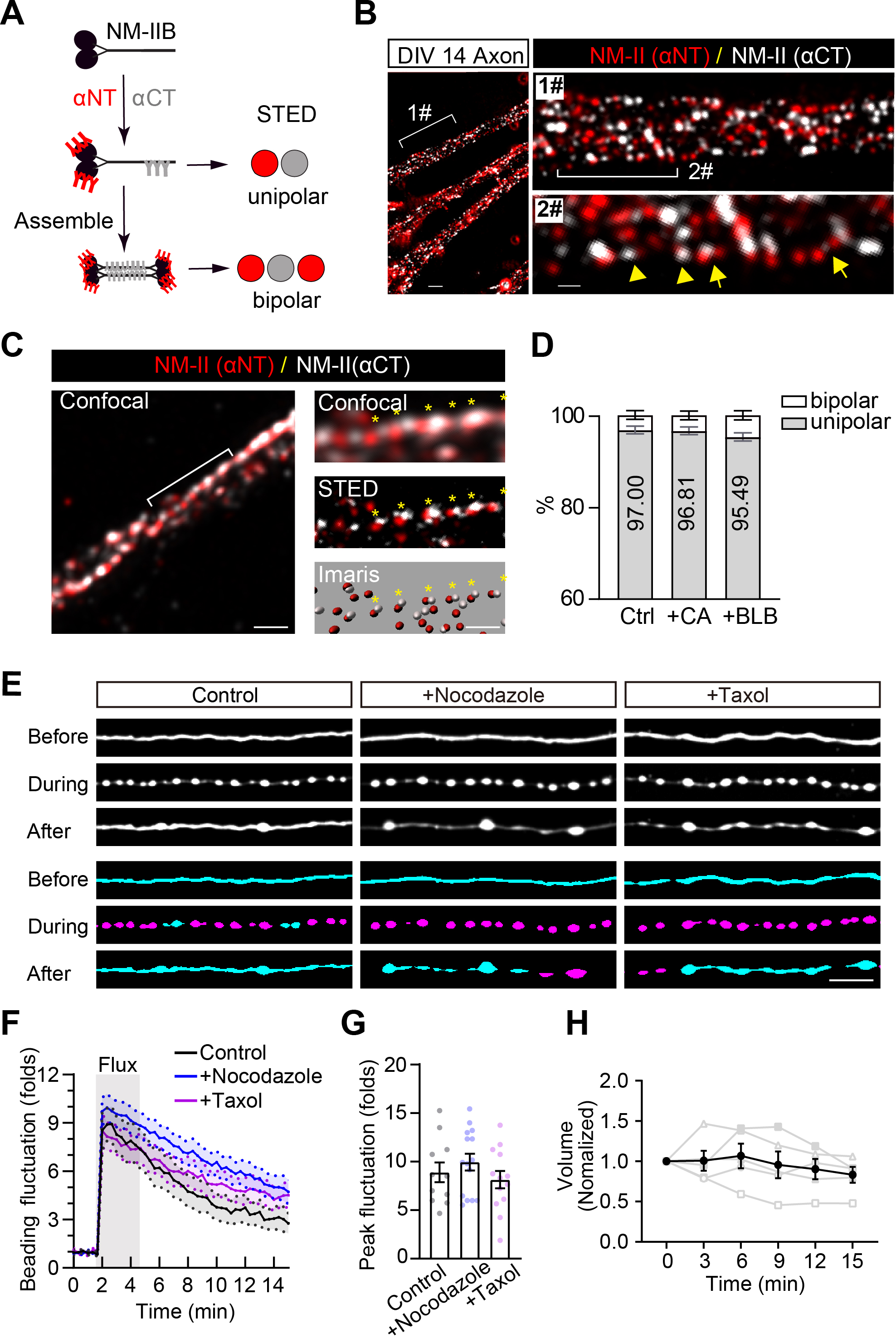
Reversible axon beading is not caused by the disruption of microtubule tracks. **(a)** Cartoon showing the unipolar or bipolar NM-II filaments represented by double or triple-dot immunostaining signals revealed by antibodies recognizing the C-terminal (grey) and N-terminal (red) of the NM-IIB heavy chain. **(b)** STED image showing the distribution of the NM-II filaments beneath the plasm membrane of the axon. The unipolar and bipolar NM-II filaments are indicated with arrowheads and arrows, respectively. Bar = 1 μm (left) and 300 nm (right). **(c)** The STED images were rendered into spots using the “spot” function of the Imaris software, with unipolar filaments indicated by yellow asterisks. Bar = 1 μm (left) and 0.5 μm (right). **(d)** Quantification of the ratio of unipolar or bipolar NM-II filaments in the axon shafts in untreated control neurons (Ctrl), and neurons treated with either Calyculin A (+CA; 50 nM) or Blebbistatin (+BLB; 50 μM). N = 6, 9, 9, respectively. **(e)** Rat hippocampal neurons cultured in an AoC device were transfected with Lifeact-GFP on DIV5-6. On DIV7-8, after being pre-treated with either a microtubule (MT) polymerization inhibitor (+Nocodazole; 50 μμ) or MT stabilizer (+Taxol; 10 μμ) for 30 min, neurons were subjected to 50 μL/min-flux for 180 s. Time-lapse images showing the deformation of the same axons before, during and after the flux are presented in the top panels. The automatically detected “beading” and “between” segments are shown in the bottom panels. Bar = 5 μm. **(f-h)** Quantification of the dynamic beading formation **(f)**, peak beading **(g)** and axon volume changes **(h)** before, during and after the flux are presented. N = 11, 15, 13, respectively. Results are shown as mean ± s.e.m.; (d, g) unpaired two-tailed student’s *t*-test, (h) paired two-tailed student’s *t*-test; n.s. Non-significant.

**Supplementary Figure 4.**
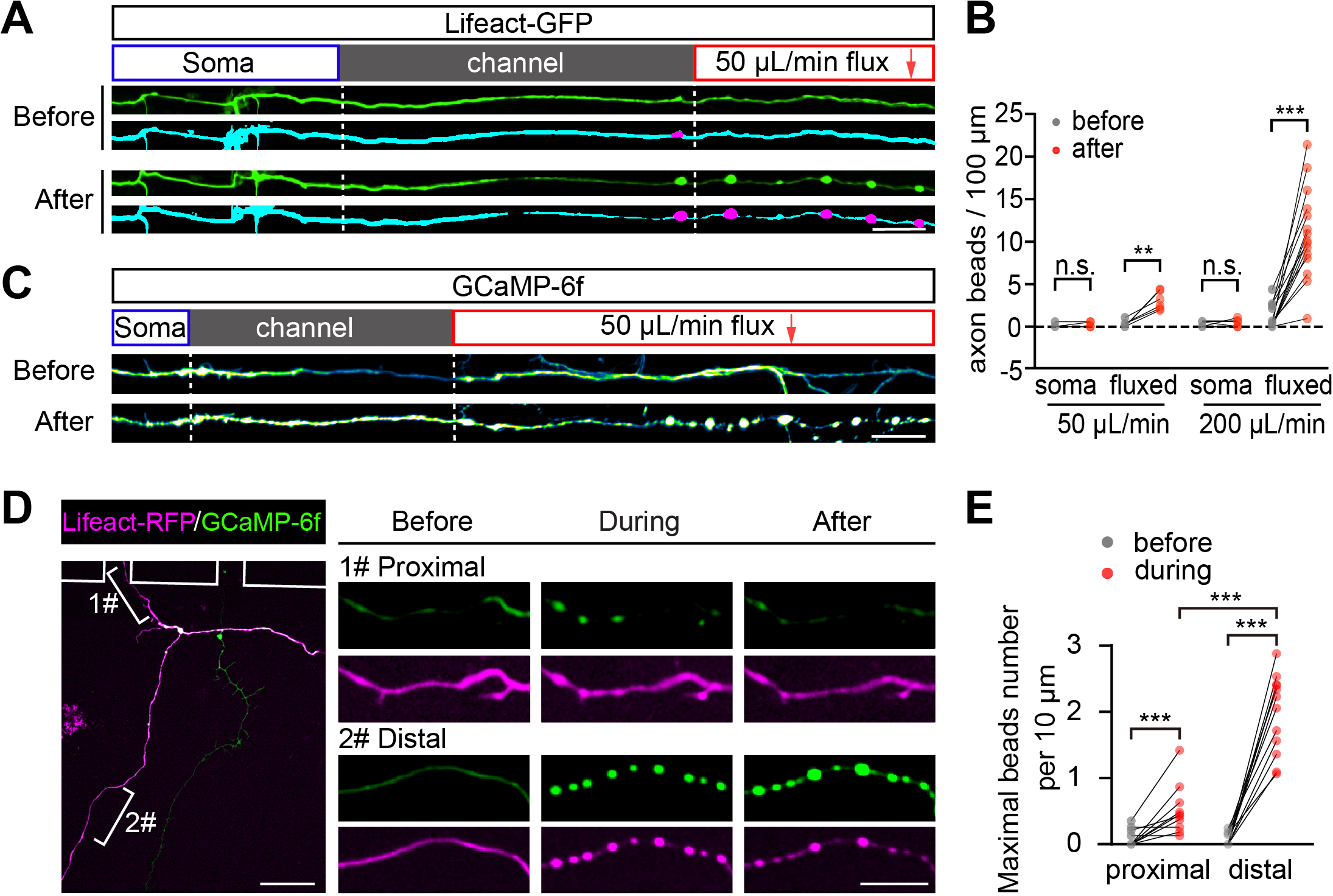
Stress-induced axonal beading and Ca^2+^ elevations are restricted to the stressed segments of axons. **(a)** Representative images of the straightened axon before and after the flux (top) and automatic detection of the axon beading (bottom). Bar = 20 μm. **(b)** Paired comparison of axon beading density in soma chamber (soma) and central injury chamber (fluxed). N = 6, 6, 21, 15, respectively. **(c)** Representative images showing the [Ca^2+^]_axon_ intensity in straightened axons before and after the flux. Bar = 20 μm. **(d)** Rat hippocampal neurons expressing Lifeact-RFP and GCaMP-6f were fluxed at 50 μL/min. Representative field showing deformation of non-stressed axonal segments near the boundary of the AoC device (#1, proximal) or the stressed distal part (#2, distal). Bar = 30 μm (left) and 10 μm (right). **(e)** Quantification of **(d)**. N = 10, 12, respectively. In (b, e) paired two-tail students’ *t*-test; \*\**p* < 0.01, \*\*\**p* < 0.001, n.s. Non-significant.

**Supplementary Figure 5.**
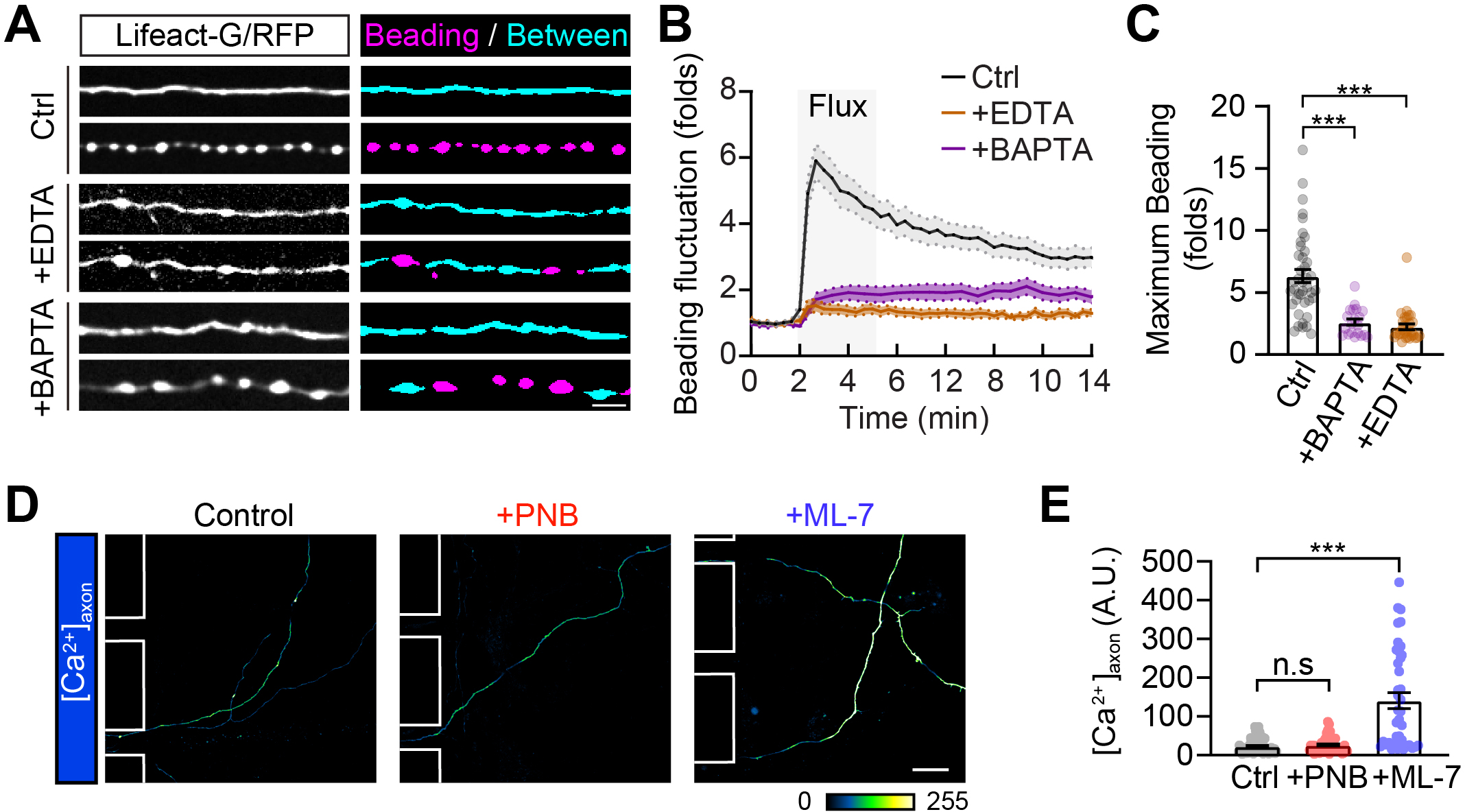
Actomyosin-II inhibitor ML-7 affects the resting Ca^2+^ level in treated axons. **(a)** Rat hippocampal neurons cultured in the AoC devices were transfected with Lifeact-GFP on DIV5-6. On DIV7-8, neurons in the area of interest were subjected to 50 μL/min flux for 180 s after being pre-treated with Ca^2+^ chelators EDTA (0.5 mμ) and BAPTA-AM (10 μμ) for 30 min. Representative time-lapse images showing the deformation before and during the flux are presented in the left panels, with the automatically detected “beading” and “between” segments shown in the right panels. Bar = 5 μm. **(b)** Quantification of the axon beading in **(a). (c)** Quantification of maximal axon beading density induced by the 50 μL/min flux for 180 s. N = 41, 29, 49, respectively**. (d)** Representative images showing the resting [Ca^2+^]_axon_ intensity before the flux in control, PNB (50 μμ) and ML-7 (10 μμ) treated axons, respectively. Bar=20 μm. **(e)** Quantification of **(d)**. N= 64, 40, 49 for control, PNB and ML-7 groups, respectively. Results are shown as mean ± s.e.m.; In (c, e) unpaired two-tailed student’s *t*-test; \*\*\**p*<0.001, n.s. Non-significant.

## Abbreviations

CNS: Central nervous system
TBI: Traumatic brain injury
DAI: Diffuse axonal injury
MT: Microtubule
MPS: Membrane-associated periodic cytoskeleton
NM-II: Non-muscle myosin II
STED: Stimulated Emission Depletion
AoC: Axon-on-a-chip
DIV: Days *in vitro*
IF: Immunofluorescent
SIM: Structured Illumination Microscope
SEM: Scanning Electron Microscope
FAS: Focal axonal swellings
TEM: Transmission Electronic Microscope
BLB: Blebbistatin
CA: Calyculin A
LatB: Latrunculin B
MRLC: Myosin regulatory light chain
p-MLC: the phosphorylation of MRLC
PNB: Para-Nitro-Blebbistatin
AAV: Adeno-associated virus

## Notes

### Competing Interest Statement

The authors have declared no competing interest.

